# Inhibitory muscarinic acetylcholine receptors enhance aversive olfactory conditioning in adult Drosophila

**DOI:** 10.1101/382440

**Authors:** Noa Bielopolski, Hoger Amin, Anthi A. Apostolopoulou, Eyal Rozenfeld, Hadas Lerner, Wolf Huetteroth, Andrew C. Lin, Moshe Parnas

## Abstract

Olfactory associative learning in *Drosophila* is mediated by synaptic plasticity between the Kenyon cells of the mushroom body and their output neurons. Both Kenyon cells and their inputs are cholinergic, yet little is known about the physiological function of muscarinic acetylcholine receptors in learning in adult flies. Here we show that aversive olfactory learning in adult flies requires type A muscarinic acetylcholine receptors (mAChR-A) specifically in the gamma subtype of Kenyon cells. Surprisingly, mAChR-A inhibits odor responses in both Kenyon cell dendrites and axons. Moreover, mAChR-A knockdown impairs the learning-associated depression of odor responses in a mushroom body output neuron. Our results suggest that mAChR-A is required at Kenyon cell presynaptic terminals to depress the synapses between Kenyon cells and their output neurons, and may suggest a role for the recently discovered axo-axonal synapses between Kenyon cells.

## Introduction

Animals learn to modify their behavior based on past experience by changing connection strengths between neurons, and this synaptic plasticity often occurs through metabotropic receptors. In particular, neurons commonly express both ionotropic and metabotropic receptors for the same neurotransmitter, where the two may mediate different functions (e.g., direct excitation/inhibition vs. synaptic plasticity). In mammals, where glutamate is the principal excitatory neurotransmitter, metabotropic glutamate receptors (mGluRs) have been widely implicated in synaptic plasticity and memory (Jörntell and Hansel, 2006; Lüscher and Huber, 2010). Given the complexity of linking behavior to artificially induced plasticity in brain slices (Schonewille et al., 2011; Yamaguchi et al., 2016), it would be useful to study the role of metabotropic receptors in learning in a simpler genetic model system with a clearer behavioral readout of synaptic plasticity. One such system is *Drosophila,* where powerful genetic tools and well-defined anatomy have yielded a detailed understanding of the circuit and molecular mechanisms underlying associative memory (Busto et al., 2010; Cognigni et al., 2017; Hige, 2017). The principal excitatory neurotransmitter in *Drosophila* is acetylcholine, but, surprisingly, little is known about the function of metabotropic acetylcholine signaling in synaptic plasticity or neuromodulation in *Drosophila*. Here we address this question using olfactory associative memory.

Flies can learn to associate an odor (conditioned stimulus, CS) with a positive (sugar) or a negative (electric shock) unconditioned stimulus (US), so that they later approach ‘rewarded’ odors and avoid ‘punished’ odors. This association is thought to be formed in the presynaptic terminals of the ~2,000 Kenyon cells (KCs) that make up the mushroom body (MB), the fly’s olfactory memory center (Busto et al., 2010; Cognigni et al., 2017; Hige, 2017). These KCs are activated by odors via second-order olfactory neurons called projection neurons (PNs). Each odor elicits responses in a sparse subset of KCs (Campbell et al., 2013; Lin et al., 2014) so that odor identity is encoded in which KCs respond to each odor. When an odor (CS) is paired with reward/punishment (US), an odor-specific set of KCs is activated at the same time that dopaminergic neurons (DANs) release dopamine onto KC presynaptic terminals. The coincident activation causes long-term depression (LTD) of synapses from the odor-activated KCs onto mushroom body output neurons (MBONs) that lead to approach or avoidance behavior (Aso and Rubin, 2016; Aso et al., 2014b; Cohn et al., 2015; Hige et al., 2015b; Owald et al., 2015; Perisse et al., 2016; Séjourné et al., 2011). In particular, training specifically depresses KC-MBON synapses of the ‘wrong’ valence (e.g., odor-punishment pairing depresses odor responses of MBONs that lead to approach behavior), because different pairs of ‘matching’ DANs/MBONs (e.g. punishment/approach, reward/avoidance) innervate distinct regions along KC axons (Aso et al., 2014a). Yet recent studies show that the mushroom body contains not just KC->MBON and DAN->KC synapses, but also KC->DAN, DAN->MBON and even KC->KC synapses (Cervantes-Sandoval et al., 2017; Eichler et al., 2017; Takemura et al., 2017), suggesting that this model may be incomplete.

Both MB input (PNs) and output (KCs) are cholinergic (Barnstedt et al., 2016; Yasuyama and Salvaterra, 1999), and KCs express both ionotropic (nicotinic) and metabotropic (muscarinic) acetylcholine receptors (Croset et al., 2018; Davie et al., 2018). The nicotinic receptors mediate fast excitatory synaptic currents (Su and O’Dowd, 2003), while the physiological function of the muscarinic receptors is unknown. Muscarinic acetylcholine receptors (mAChRs) are G-protein coupled receptors; out of the three mAChRs in *Drosophila* (mAChR-A, mAChR-B and mAChR-C), mAChR-A (also called Dm1, mAcR-60C or mAChR) is the most closely homologous to mammalian mAChRs (Collin et al., 2013). Mammalian mAChRs are typically divided between ‘M_1_-type’ (M_1_/M_3_/M_5_), which signal via G_q_ and are generally excitatory, and ‘M_2_-type’ (M_2_/M_4_), which signal via G_i/0_ and are generally inhibitory (Caulfield and Birdsall, 1998). *Drosophila* mAChR-A seems to use ‘M_1_-type’ signaling: when heterologously expressed in Chinese hamster ovary (CHO) cells, it signals via G_q_ protein (Collin et al., 2013; Ren et al., 2015) to activate phospholipase C, which produces inositol trisphosphate to release Ca^2+^ from internal stores.

Recent work indicates that mAChR-A is required for aversive olfactory learning in *Drosophila* larvae, as knocking down mAChR-A expression in KCs impairs learning (Silva et al., 2015). However, it is unclear whether mAChR-A is involved in olfactory learning in adult *Drosophila,* given that mAChR-A is thought to signal through G_q_, and in adult flies G_q_ signaling downstream of the dopamine receptor Damb promotes forgetting, not learning (Berry et al., 2012; Himmelreich et al., 2017). Moreover, it is unknown how mAChR-A affects the activity or physiology of KCs, where it acts (at KC axons or dendrites or both), and how these effects contribute to olfactory learning.

Here we show that mAChR-A is required in KCs for aversive olfactory learning in adult *Drosophila*. Surprisingly, genetic and pharmacological manipulations of mAChR-A suggest that mAChR-A is inhibitory and acts in part on KC axons. Moreover, mAChR-A knockdown impairs the learning-associated depression of odor responses in a key MB output neuron, MB-MVP2. We suggest that mAChR-A is required to depress synapses between KCs and their outputs.

## Results

### mAChR-A expression in KCs is required for aversive olfactory learning in adult flies

*Drosophila* larvae with reduced mAChR-A expression in KCs show impaired aversive olfactory learning (Silva et al., 2015), but it remains unknown whether mAChRA in KCs also functions in learning in adult flies. We addressed this question by knocking down mAChR-A expression in KCs using two UAS-RNAi lines, “RNAi 1” and “RNAi 2” (see Methods). Only RNAi 2 requires co-expression of Dicer-2 (Dcr-2) for optimal knockdown. To test the efficiency of these RNAi constructs, we expressed them pan-neuronally using elav-GAL4 and measured their effects on mAChR-A expression levels using quantitative real time polymerase chain reaction (qRT-PCR). Both RNAi lines strongly reduce mAChR-A levels (RNAi 1: 36±4% of elav-GAL4 control, i.e., 64±4% below normal; RNAi 2: 34±2% of normal; mean±s.e.m.; see **Figure 1A**). We then examined whether knocking down mAChR-A in KCs using the pan-KC driver OK107-GAL4 affects short term aversive conditioning in adult flies. We used the standard training protocol and odors used in the field (i.e. 3-octanol, OCT, and 4-methylcyclohexanol, MCH; see Methods). Under these conditions both UAS-RNAi transgenes significantly reduced aversive conditioning (**Figure 1B,C** and **Figure S1A**). Knocking down mAChR-A had no effect on naive avoidance of MCH and OCT (**Figure 1D**; see Methods) or flies’ reaction to electric shock (**Figure S1B**), showing that the defect was specific to learning, rather than reflecting a failure to detect odors or shock.

**Figure 1:**
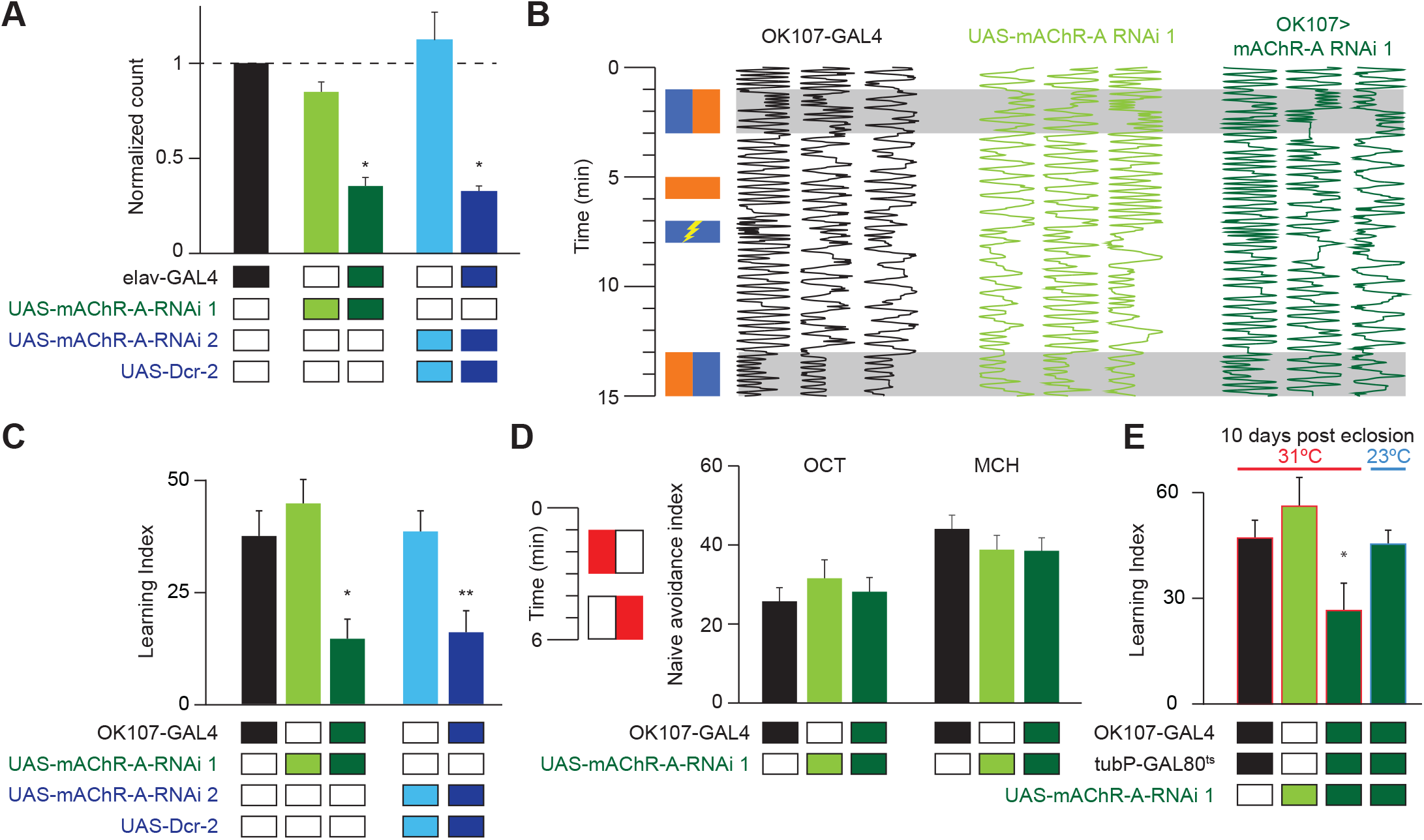
mAChR-A is required in the MB for short term aversive olfactory learning and memory but not for naïve behavior. **(A)** qRT-PCR of mAChR-A with mAChR-A RNAi driven by elav-GAL4. The housekeeping gene eEF1α2 (eukaryotic translation elongation factor 1 alpha 2, CG1873) was used for normalization. Knockdown flies have ~30% of the control levels of mAChR-A mRNA (mean ± SEM; number of biological replicates (left to right): 6, 7, 7, 4, 4, each with 3 technical replicates; * p < 0.05; Kruskal-Wallis test with Dunn’s multiple comparisons test). For detailed statistical analysis see **Table S1**. **(B)** Each trace shows the movement of an individual fly during the training protocol, with fly position in the chamber (horizontal dimension) plotted against time (vertical dimension). Colored rectangles illustrate which odor is presented on each side of the chamber during training and testing. Flies were conditioned against MCH (blue rectangles; see Methods). **(C)** Learning scores in flies with mAChR-A RNAi driven by OK107-GAL4. mAChR-A knockdown reduced learning scores compared to controls (mean ± SEM, n (left to right): 69, 69, 70, 71, 71, * p < 0.05, ** p<0.01; Kruskal-Wallis test with Dunn’s multiple comparisons test). **(D)** mAChR-A KD flies show normal olfactory avoidance to OCT and MCH compared to their genotypic controls (mean ± SEM, n (left to right): 68, 67, 58, 63, 91, 67, p = 0.82 for OCT, p = 0.64 for MCH; Kruskal-Wallis test). Colored rectangles show stimulus protocol as in (B); red for odor (MCH or OCT), white for air. **(E)** Learning scores in flies with mAChR-A RNAi 1 driven by OK107-GAL4 with GAL80^ts^ repression. Flies raised at 23 °C and heated to 31 °C as adults (red outlines) had impaired learning compared to controls. Control flies kept at 23 °C throughout (blue outline), thus blocking mAChR-A RNAi expression, showed no learning defects (mean ± SEM, n (left to right): 51, 41, 58, 51, ** p < 0.05, Kruskal-Wallis test with Dunn’s multiple comparisons test). For detailed statistical analysis see Table S1.

Given that mAChR-A is expressed in the larval MB and indeed contributes to aversive conditioning in larvae, it is possible that developmental effects underlie the reduced learning observed in mAChR-A KD flies. To test this, we used tub-GAL80^ts^ to suppress RNAi 1 expression during development. Flies were grown at 23°C until 3 days after eclosion and were then transferred to 31 °C for 7 days. Adult-only knockdown of mAChR-A in KCs reduced learning (**Figure 1E**), just as constitutive knockdown did, indicating that mAChR-A plays a physiological, not purely developmental, role in aversive conditioning. To further verify that GAL80^ts^ efficiently blocks RNAi expression (i.e., that GAL80^ts^ is not leaky), flies were grown at 23°C without transferring them to 31°C, thus blocking RNAi expression also in adults. When tested for learning at 10 days old, these flies showed normal learning (**Figure 1E**).

### mAChR-A is required for olfactory learning in γ KCs, not αβ or α′β′ KCs

Kenyon cells are subdivided into three main classes: γ neurons project to the horizontal lobes only, while the axons of αβ and α′β′ neurons bifurcate to form the α and α′ portions of the vertical lobes and the β and β′ portions of the horizontal lobes. These different classes play different roles in olfactory conditioning (Guven-Ozkan and Davis, 2014; Krashes et al., 2007). To unravel in which class(es) mAChR-A functions, we used a Minos-mediated integration cassette (MiMIC) line to investigate where mAChR-A is expressed (Venken et al., 2011). The MiMIC insertion in mAChR-A lies in the first 5’ non-coding intron, creating a gene trap where GFP in the MiMIC cassette should be expressed in whichever cells endogenously express mAChR-A. Because the GFP in the original mAChR-A MiMIC cassette produced very little fluorescent signal (data not shown), we used recombinase-mediated cassette exchange (RMCE) to replace the original MiMIC cassette with a MiMIC cassette containing GAL4 (Venken et al., 2011). These new mAChR-A-MiMIC-GAL4 flies should express GAL4 wherever mAChR-A is endogenously expressed. To reveal the expression pattern of mAChR-A, we crossed mAChR-A-MiMIC-GAL4 and 20xUAS-eGFP flies. mAChR-A-MiMIC-GAL4 drove GFP expression throughout the brain, consistent with previous reports (Blake et al., 1993; Croset et al., 2018; Davie et al., 2018; Hannan and Hall, 1996) and with the fact that the *Drosophila* brain is mostly cholinergic. In the mushroom bodies, GFP was expressed in the αβ and γ lobes, but not the α′β′ lobes (**Figure 2A**). No GFP signal was observed with an inverted insertion where GAL4 is inserted in the MiMIC locus in the wrong direction (data not shown). Consistent with these MiMIC results, two recently reported databases of single-cell transcriptomic analysis of the *Drosophila* brain (Croset et al., 2018; Davie et al., 2018) confirm that mAChR-A is more highly expressed in αβ and γ KCs than in α′β′ KCs (**Figure S2**).

**Figure 2:**
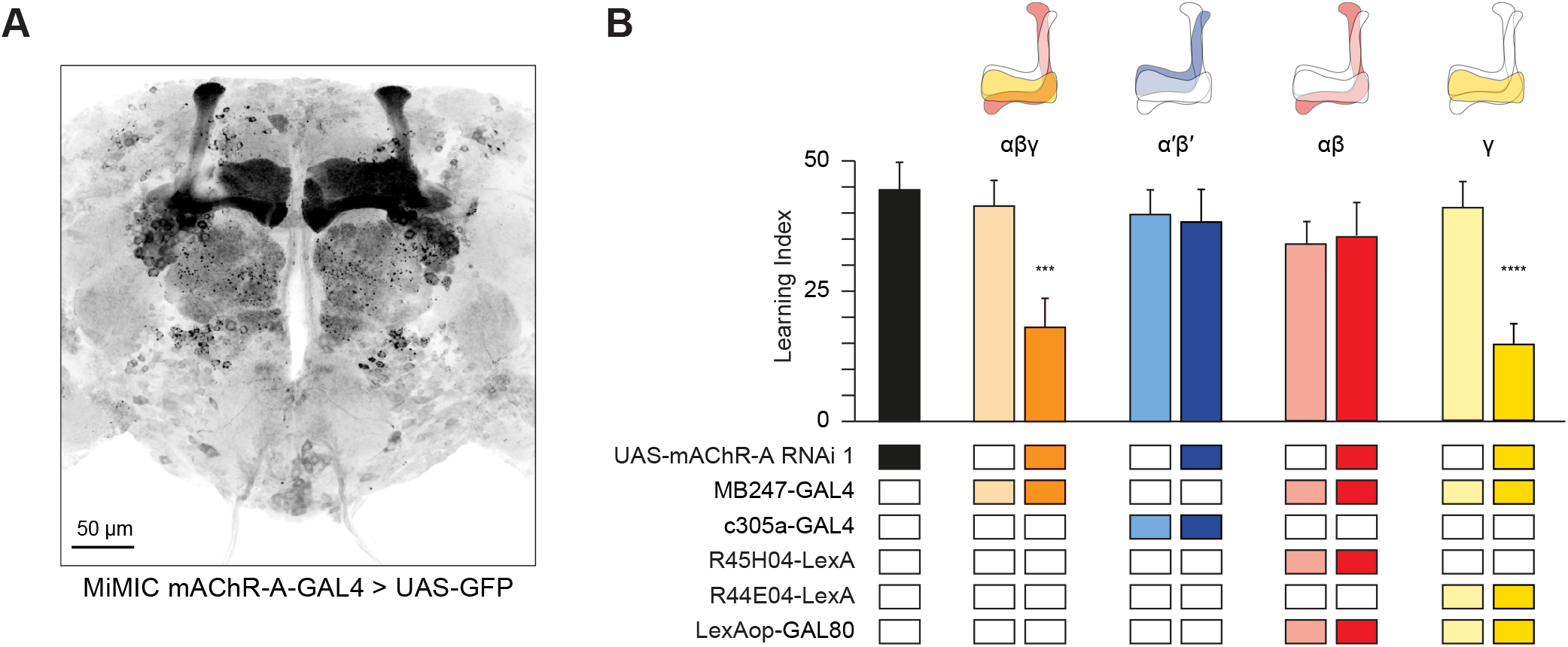
mAChR-A is required for short term aversive olfactory learning and memory in γ KCs. **(A)** Maximum intensity projection of 70 confocal sections (2 μm) through the central brain of a fly carrying MiMIC-mAChR-A-GAL4 and 20xUAS-6xGFP transgenes. MB αβ and γ lobes are clearly observed. No GFP expression is observed in α′β′ lobes. **(B)** mAChR-A RNAi 1 was targeted to different subpopulations of KCs. Learning scores were reduced compared to controls when mAChR-A RNAi 1 was expressed in αβ and γ KCs or γ KCs alone, but not when mAChR-A RNAi 1 was expressed in αβ or α′β′ KCs. (mean ± SEM, n (left to right): 69, 51, 70, 76, 69, 60, 68, 66, 71, *** p < 0.001, Kruskal-Wallis test with Dunn’s multiple comparisons test). For detailed statistical analysis see **Table S1**. The data for the UAS-mAChR-A RNAi 1 control are duplicated from **Figure 1**.

The higher expression of mAChR-A in αβ and γ KCs compared to α′β′ KCs suggests that learning would be impaired by mAChR-A knockdown in αβ or γ, but not α′β′, KCs. To test this, we expressed mAChR-A RNAi in different KC classes. As expected, aversive olfactory learning was reduced by knocking down mAChR-A in αβ and γ KCs together using MB247-GAL4, but not by knockdown in α′β′ KCs using c305a-GAL4. To examine if αβ and γ KCs both participate in the reduced learning observed in mAChR-A knockdown flies, we sought to limit mAChR-A RNAi expression to either αβ or γ neurons. While strong driver lines exist for αβ neurons, the γ GAL4 drivers we tested were fairly weak (H24-GAL4, MB131B, R45H04-GAL4, data not shown), perhaps too weak to drive mAChR-A-RNAi enough to knock down mAChR-A efficiently. Therefore, we used MB247-GAL4, which was strong enough to affect behavior, and blocked GAL4 activity in either αβ or γ KCs by expressing the GAL80 repressor under the control of R44E04-LexA (αβ KCs) or R45H04-LexA (γ KCs) (Bräcker et al., 2013). These combinations drove strong, specific expression in αβ or γ KCs (**Figure S3**). Learning was reduced by mAChR-A RNAi expression in γ, but not αβ, KCs (**Figure 2B**). These results suggest that mAChR-A is specifically required in γ KCs for aversive olfactory learning and short-term memory.

### mAChR-A suppresses odor responses in γ KCs

We next asked what effect mAChR-A knockdown has on the physiology of KCs, by expressing GCaMP6f and mAChR-A RNAi 2 together in KCs using OK107-GAL4. Knocking down mAChR-A in KCs increased odor-evoked Ca^2+^ influx in the mushroom body calyx, where KC dendrites reside (**Figure 3**). This result is somewhat surprising because mAChR-A is a G_q_ coupled receptor whose activation leads to Ca^2+^ release from internal stores (Ren et al., 2015), which predicts that mAChR-A knockdown should decrease, not increase, odor-evoked Ca^2+^ influx in KCs. However, some examples have been reported of inhibitory signaling through G_q_ by M_1_-type mAChRs (see Discussion), and *Drosophila* mAChR-A may join these as another example of an inhibitory mAChR signaling through Gq.

**Figure 3:**
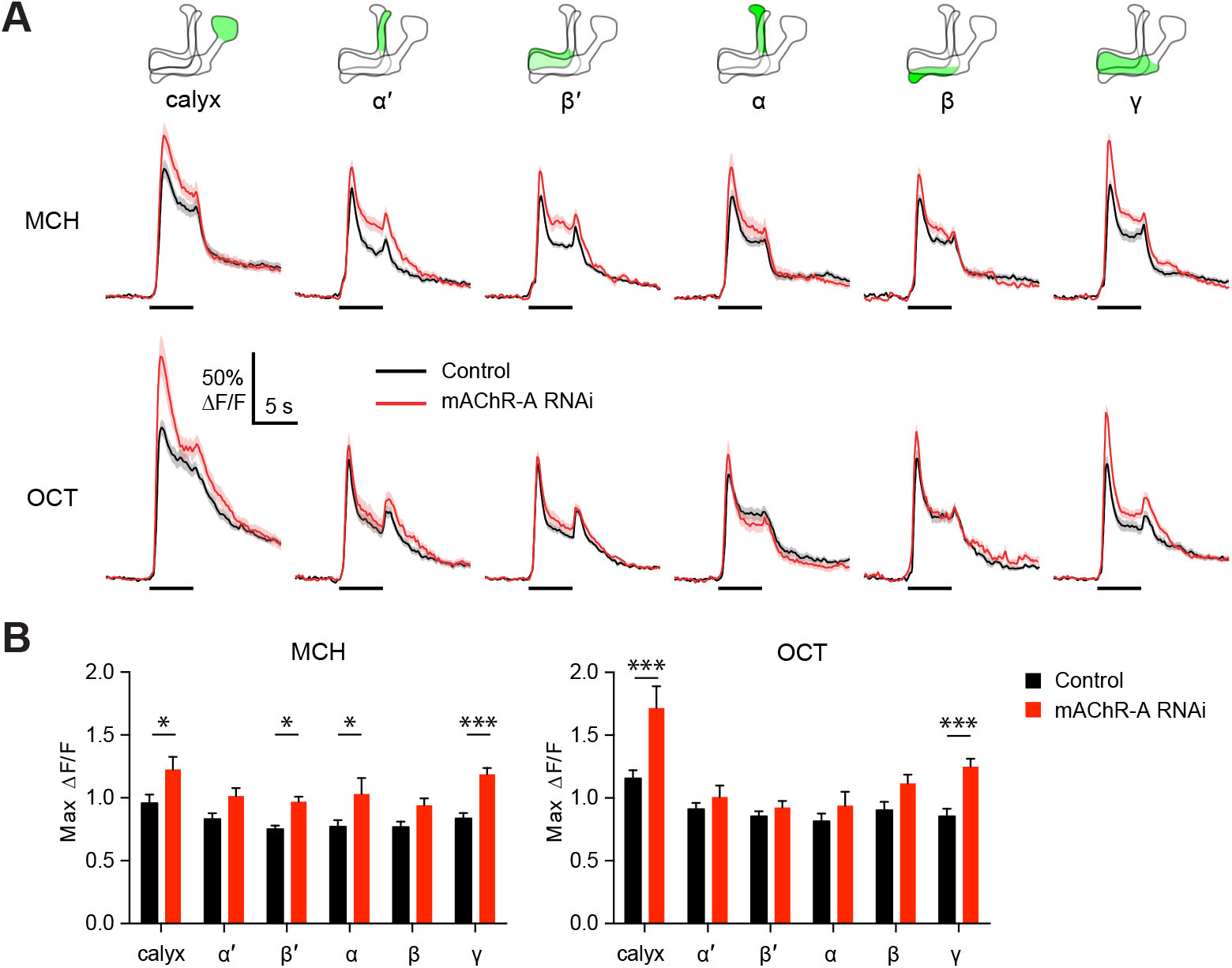
mAChR-A knockdown increases odor responses in γ KCs. Odor responses to MCH and OCT were measured in control (OK107-GAL4>GCaMP6f, Dcr-2) and knockdown (OK107-GAL4>GCaMP6f, Dcr-2, mAChR-A-RNAi 2) flies. **(A)** ΔF/F of GCaMP6f signal in different areas of the MB in control (black) and knockdown (red) flies, during presentation of odor pulses (horizontal lines). Data are mean (solid line) ± SEM (shaded area). Diagrams illustrate which region of the MB was analyzed. **(B)** Peak response of the traces presented in A (mean ± SEM.) n given as number of hemispheres (number of flies) for control and knockdown flies, respectively: calyx, 23 (13), 17 (10); α and α′, 24 (13), 20 (10); β, β′ and γ, 27 (14), 22 (11). * p < 0.05, *** p < 0.001, 2-way ANOVA with Holm-Sidak multiple comparisons test). For detailed statistical analysis see **Table S1**.

Because mAChR-A is required for aversive learning in γ KCs, not αβ or α′β′ KCs (**Figure 2**), we next asked if odor responses in αβ, α′β′ and γ KCs are differentially affected by mAChR-A knockdown. αβ, α′β′ and γ KC dendrites are not clearly segregated in the calyx, so we examined odor responses in the axonal lobes. Indeed, although odor responses in all lobes were increased by mAChR-A knockdown, only in the γ lobe was the effect statistically significant for both MCH and OCT (**Figure 3**). This result is consistent with the behavioral requirement for mAChR-A only in γ KCs. However, we do not rule out the possibility that mAChR-A knockdown affects αβ and α′β′ odor responses in a more subtle way that does not affect short-term memory, especially as αβ and α′β′ odor responses were somewhat, though non-significantly, increased.

Do increased odor responses in γ KCs prevent learning by increasing the overlap between the γ KC population representations of the two odors used in our task (Lin et al., 2014)? When GCaMP6f and mAChR-A-RNAi 2 were expressed in all KCs, mAChRA knockdown did not affect the sparseness or inter-odor correlation of KC population odor responses (**Figure 4A-C**) even though it increased overall calyx responses. To focus specifically on γ KCs, we expressed GCaMP6f and mAChR-A-RNAi 1 only in γ KCs, using mb247-Gal4, R44E04-LexA and lexAop-GAL80, as in **Figure 2B**. GCaMP6f was visible mainly in the γ lobe (**Figure 4D**). γ-only expression of mAChR-A-RNAi 1 increased odor responses in the calyx (here, dendrites of γ KCs only) and, in the case of OCT, in the γ lobe (**Figure 4E,F**). Note that γ KC odor responses are increased by both RNAi 1 (**Figure 3A,B**) and RNAi 2 (**Figure 4E,F**). As with pan-KC expression, γ-only expression of mAChR-A-RNAi 1 did not affect the sparseness or inter-odor correlation of γ KCs (**Figure 4G-I**). Thus, mAChR-A knockdown does not impair learning through increased overlap in KC population odor representations.

**Figure 4:**
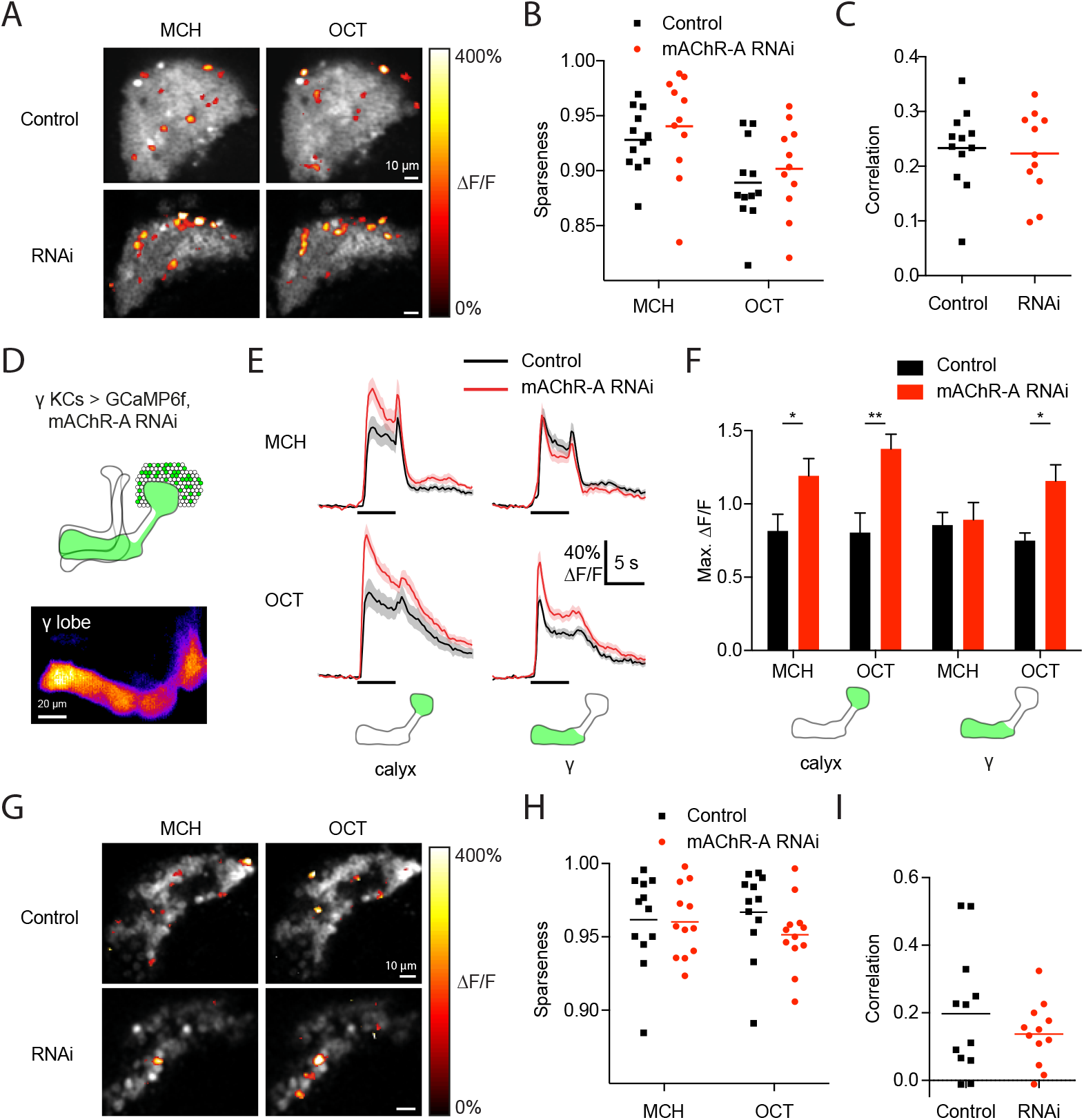
mAChR-A knockdown does not affect KC odor identity coding. **(A)** Example activity maps (single optical sections from a z-stack) of KC odor responses to MCH and OCT in control (OK107-GAL4>GCaMP6f, Dcr-2) and mAChR-A knockdown (OK107-GAL4>GCaMP6f, Dcr-2, mAChR-A-RNAi 2) flies where all KCs are imaged. False-coloring indicates ΔF/F of the odor response, overlaid on grayscale baseline GCaMP6f signal. Scale bar, 10 μm. For detailed statistical analysis see **Table S1**. **(B)** Sparseness of pan-KC population responses is not affected by mAChR-A knockdown (p = 0.38, 2-way repeated-measures ANOVA). **(C)** Correlation between pan-KC population responses to MCH and OCT is not affected by mAChR-A knockdown (p = 0.75, t-test). **(D)** Upper: diagram of γ KCs (green). Lower: False-coloured average-intensity Z-projection of the horizontal lobe in a control fly imaged from a dorsal view in panel E (mb247-GAL4>GCaMP6f, R44E04-LexA>GAL80), averaged over 10 s before the odor stimulus. R44E04-LexA>GAL80 almost completely suppresses β lobe expression. Scale bar, 20 μm. **(E)** Knocking down mAChR-A only in γ KCs increases γ KC odor responses. Shown here are odor responses in the calyx and γ lobe of control (mb247-GAL4>GCaMP6f, R44E04-LexA>GAL80) and knockdown (mb247-GAL4>GCaMP6f, mAChR-A-RNAi 1, R44E04-LexA>GAL80) flies. **(F)** Peak response of the traces presented in D (mean ± SEM.) n given as number of hemispheres (number of flies): 11 (6) for control, 12 (6) for knockdown. * p < 0.05, ** p < 0.01, 2-way repeated-measures ANOVA with Holm-Sidak multiple comparisons test. **(G)** Example activity maps (single optical sections from a z-stack) of γ KC odor responses to MCH and OCT in control (mb247-GAL4>GCaMP6f, R44E04-LexA>GAL80) and knockdown (mb247-GAL4>GCaMP6f, mAChR-A-RNAi 1, R44E04-LexA>GAL80) flies. Note the gaps in baseline GCaMP6f signal due to lack of αβ and α′β′ KCs labeled. Scale bar, 10 μm **(H)** Sparseness of γ KC population responses is not affected by mAChR-A knockdown (p = 0.76, 2-way repeated-measures ANOVA). **(I)** Correlation between γ KC population responses to MCH and OCT is not affected by mAChR-A knockdown (p = 0.32, t-test).

### KC odor responses are decreased by an mAChR agonist

RNAi-based knockdown of mAChR-A might induce homeostatic compensation that obscures or even reverses the primary effect of reduced mAChR-A expression. To test the acute role of mAChR-A in regulating KC activity, we took the complementary approach of pharmacologically activating mAChR-A. Initially we bath-applied 10 μM muscarine, an mAChR-A agonist *(Drosophila* mAChR-B is 1000-fold less sensitive to muscarine than mAChR-A is (Collin et al., 2013), and mAChR-C is not expressed in the brain (Davie et al., 2018)). Muscarine strongly decreased odor responses in all subtypes of KCs (**Figure 5A,B**). However, muscarine did not significantly affect odor responses in PNs (**Figure 5C**), suggesting that the effect of muscarine on KCs arose in KCs, not earlier in the olfactory pathway. KCs can be silenced by an inhibitory GABAergic neuron called the anterior paired lateral (APL) neuron (Lin et al., 2014; Masuda-Nakagawa et al., 2014; Papadopoulou et al., 2011), so we asked whether muscarine reduces KC odor responses indirectly by activating APL, rather than directly inhibiting KCs. We applied muscarine to flies with APL-specific expression of tetanus toxin (TNT), which blocks inhibition from APL and thereby greatly increases KC odor responses. In these flies, APL is labeled stochastically, so hemispheres where APL was unlabeled served as controls (Lin et al., 2014) (see Methods). Muscarine decreased KC odor responses both in control hemispheres and hemispheres where APL synaptic output was blocked by tetanus toxin (**Figure 5D**), and the effect of muscarine was not significantly different between the two cases (**Figure 5E**). This result indicates that muscarine does not act solely by activating APL or by enhancing inhibition on KCs (e.g., increasing membrane localization of GABA_A_ receptors).

**Figure 5:**
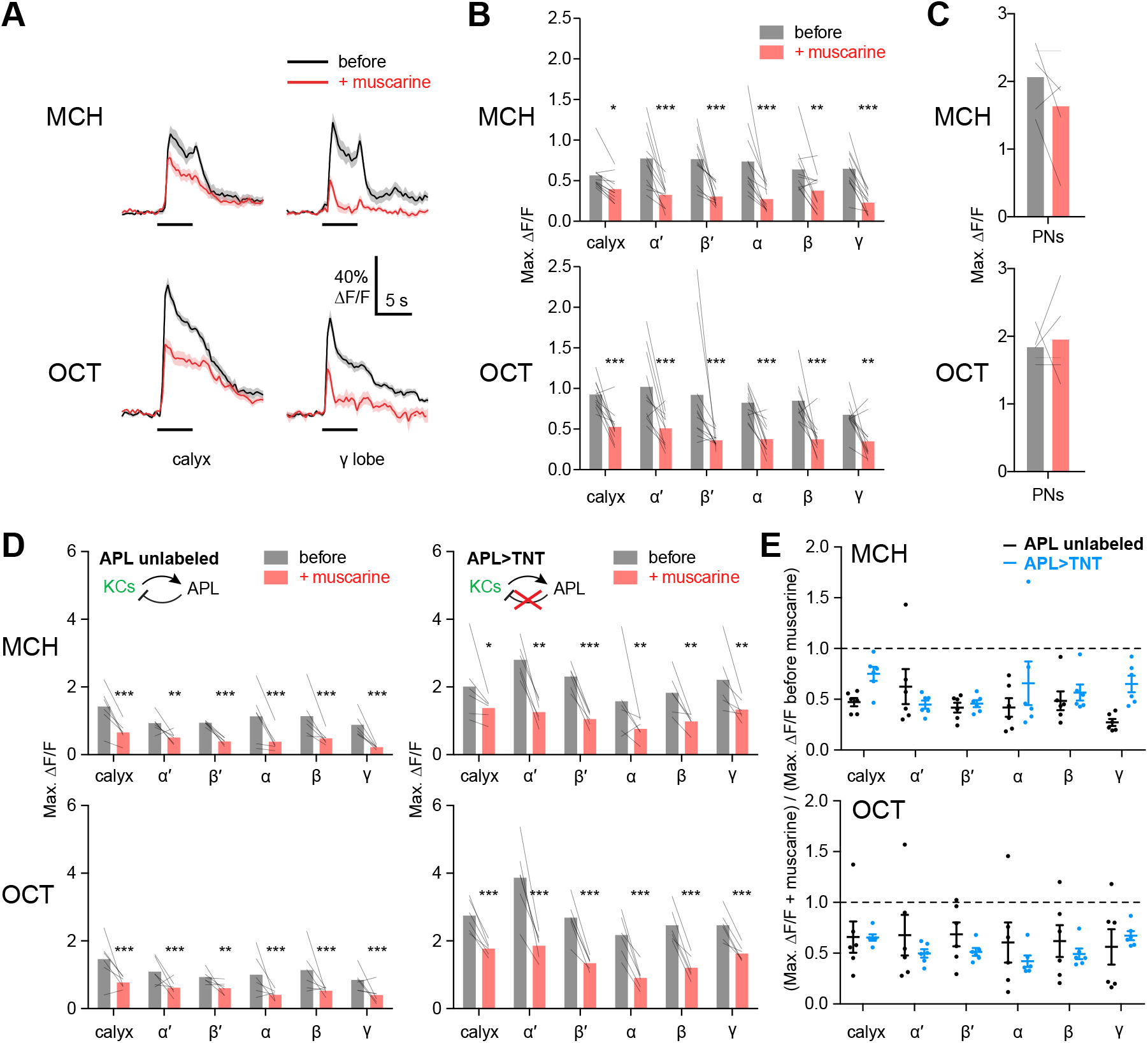
KC odor responses are decreased by muscarine. **(A)** Odor responses in the calyx and γ lobe of OK107-GAL4>GCaMP6f flies, before (black) and after (red) adding 10 μM muscarine in the bath. Data are mean (solid line) ± SEM (shaded area); horizontal lines indicate the odor pulse. Traces for all lobes are shown in Figure S4. For detailed statistical analysis see Table S1. **(B)** Peak ΔF/F during the odor pulse before and after muscarine. n = 11 hemispheres from 6 flies. * p < 0.05, ** p < 0.01, *** p < 0.001 by 2-way repeated measures ANOVA with Holm-Sidak multiple comparisons test. **(C)** Odor responses in PN axons in the calyx are not affected by 10 μM muscarine, in GH146-GAL4>GCaMP6f flies (p > 0.49, 2-way repeated measures ANOVA, n = 5 flies). **(D)** Peak ΔF/F during the odor pulse before and after muscarine in control hemispheres where APL was unlabeled (left, n = 6 hemispheres from 6 flies) and hemispheres where APL expressed tetanus toxin (TNT) (right, n = 6 hemispheres from 5 flies). * p < 0.05, ** p < 0.01, *** p < 0.001 by 2-way repeated measures ANOVA with Holm-Sidak multiple comparisons test. **(E)** (Response (peak ΔF/F during the odor pulse) after muscarine) / (response before muscarine), using data from (D). No significant differences were observed (p > 0.05, 2-way repeated measures ANOVA with Holm-Sidak multiple comparisons test).

To further clarify whether muscarine directly affects KC axons, we locally applied muscarine to the MB horizontal lobe by pressure ejection (**Figure 6**). Red dye included in the ejected solution confirmed that the muscarine did not spread to the calyx (**Figure 6B**). In the absence of odor stimuli, locally applied muscarine on the horizontal lobe decreased GCaMP6f signal in the β and γ lobes during the period 0.5 – 1.5 s after application, suggesting that mAChR-A reduces baseline Ca^2+^ levels in some KCs (**Figure 6A,C**). Interestingly, in α′β′ KCs, local muscarine application caused a slower rise in GCaMP6f signal at 3 – 4 s after application (**Figure 6A,D**), suggesting that mAChR-A has different effects in different KCs or on different timescales. To test whether locally applied muscarine also modulates odor-evoked Ca^2+^ influx, we applied muscarine 1 s before odor onset. Locally applied muscarine decreased KC odor responses for both odors in the γ and β lobes, and for OCT in all lobes (**Figure 6F,G**). The more obvious effect with OCT might arise from OCT activating more KCs than MCH does (i.e., MCH responses were sparser: **Figure 4B, Table S1**), consistent with previous reports (Lin et al., 2014; Perisse et al., 2013). Although the red dye suggests that muscarine applied to the horizontal lobe did not spread substantially to the vertical (α and α′) lobes (**Figure 6B**), these lobes may have been affected in the case of OCT because muscarine reached KC axons at the branch point. Notably, with only one exception, locally applied muscarine had no effect on either GCaMP6f signal or odor responses on the opposite hemisphere from the site of application (**Figure 6G**); nor did muscarine applied on the horizontal lobe affect GCaMP6f signal in the calyx (**Figure 6A**). In the case of OCT, during muscarine application to the horizontal lobe, odor responses in the calyx were lower on both the same side and opposite side, but given that the opposite side was affected in no other case, this exception may reflect sensory adaptation between the ‘before’ and ‘+muscarine’ recordings, rather than an effect of muscarine *per se*. Together, these results suggest that mAChR-A acts on KC axons to inhibit local Ca^2+^ influx.

**Figure 6:**
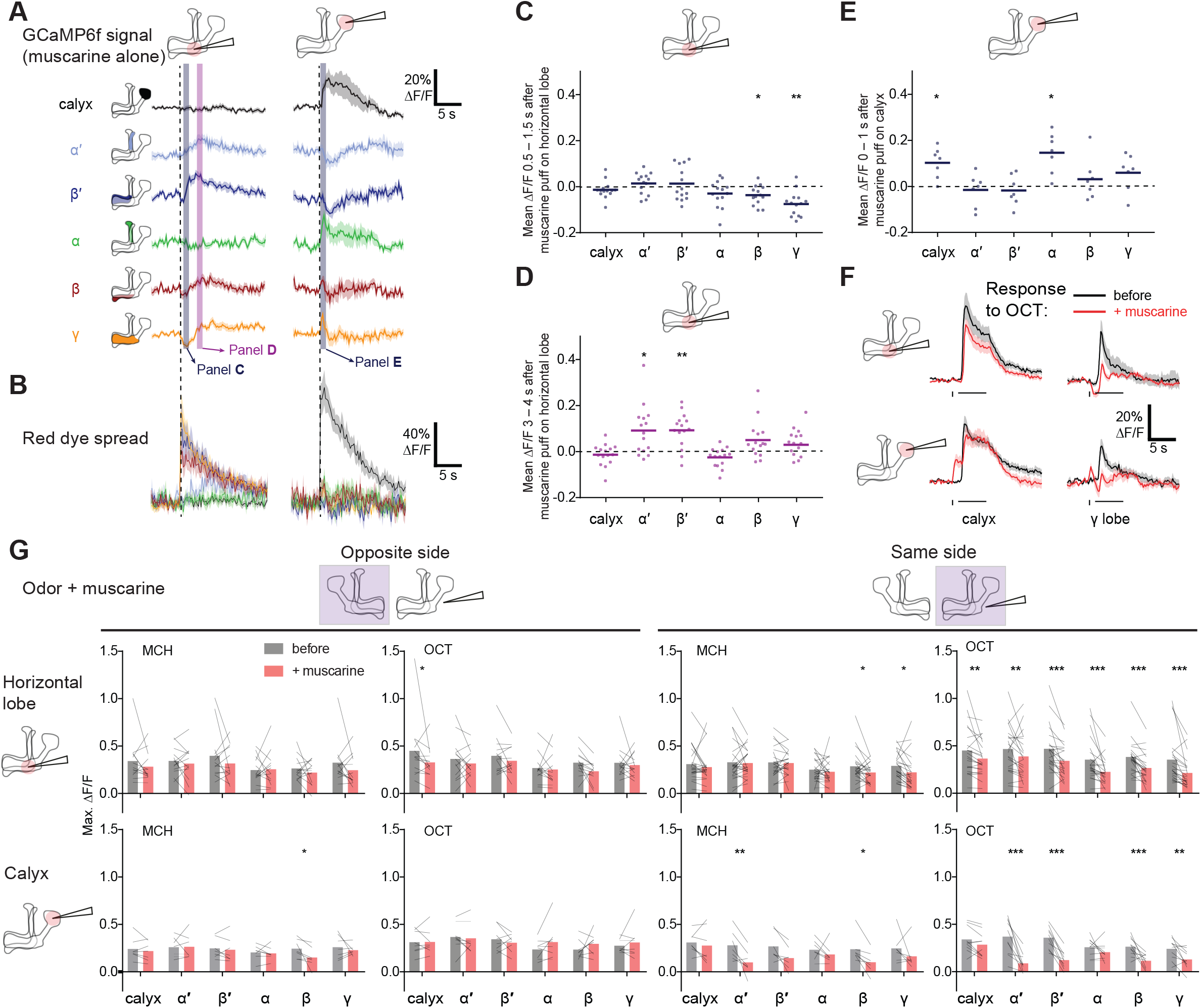
Local muscarine application to the calyx and horizontal lobe differentially affects KC subtypes. **(A)** Left: Schematic of MB, showing color scheme for the different regions where responses are quantified. Right: Average ΔF/F GCaMP6f signal in different areas of the MB of OK107>GCaMP6f flies in response to a 10 ms pulse of 20 mM muscarine on the horizontal lobe (left column) and the calyx (right column). Data are mean (solid line) ± SEM (shaded area). Dashed vertical line shows the timing of muscarine application. Gray and purple shaded bars indicate time windows used to quantify responses in panel C-E. n given as number of hemispheres (number of flies): 15 (8) for pulsing on the horizontal lobe, 7 (5) for calyx. **(B)** ΔF/F traces of red dye indicator, showing which MB regions the muscarine spread to. The traces follow the same color scheme and visuals as shown in panel A. **(C-E)** Scatter plot showing average ΔF/F of GCaMP6f signal of the different MB regions at time 0.5-1.5 s **(C)** and 3-4 s **(D)** following 10 ms pulse of 20 mM muscarine on the horizontal lobe or the calyx (E, time 0-1 s), quantified from traces shown in (A). Colors match the shaded time windows in A. n: 15 (8) for pulsing on the horizontal lobe, 7 (5) for calyx. * p < 0.05, ** p < 0.01, one-sample t-test or Wilcoxon signed-rank test (different from 0), Bonferroni correction for multiple comparisons. **(F)** Average ΔF/F GCaMP6f signal of the calyx and γ lobe during odor pulses of OCT (horizontal bar), before (black) and after (red) muscarine application on the horizontal lobe (top), or the calyx (bottom), 1 s before the odor pulse (vertical bar). Data are mean (solid line) ± SEM (shaded area). n: 12 (8) for pulsing on the horizontal lobe, 7 (5) for calyx. See **Figure S5** for all traces. **(G)** Line-bar plots showing paired peak ΔF/F GCaMP6f responses of the different MB regions during 5 s odor pulses of MCH or OCT, before (gray) and after (pink) muscarine application to the calyx or the horizontal lobe, in the hemisphere where the muscarine was applied (same side, right) or the opposite (opposite side, left). Muscarine was applied 1 s before the odor pulse. Bars show mean value. n: Horizontal lobe same side MCH 23 (15), OCT 22 (15), opposite side MCH 13 (7), OCT 13 (7). Calyx same side MCH 8 (6), OCT 10 (8), opposite side MCH 7 (5), OCT 7 (5). * p < 0.05, ** p < 0.01, *** p < 0.001 by 2-way repeated measures ANOVA with Holm-Sidak multiple comparisons test. n differs between OCT horizontal lobe, same side vs. panel **F** because a scripting error meant that for 10 of the recordings, the odor pulse paired with muscarine presentation lasted 4 s, not 5 s. These data are included in **G**, because the peak ΔF/F always occurred within the first 4 s of the odor pulse and thus was unaffected by the scripting error, but they are excluded from F because the stimuli do not match.

To test what effect mAChR-A exerts on the calyx, we locally applied muscarine to the calyx. Surprisingly, applying muscarine to the calyx in the absence of odor stimuli increased GCaMP signal in the calyx and α lobe (**Figure 6A,E**). It also decreased GCaMP signal in the α′ and β′ lobes around 1-2 s after application (**Figure 6A**), although this effect was not statistically significant. The increased Ca^2+^ in the calyx most likely did not reflect increased excitability, as applying muscarine to the calyx did not increase the calyx odor response (**Figure 6F,G**). If anything, it likely decreased the calyx odor response, because the Ca^2+^ increase induced by muscarine alone (no odor) lasted ~6-7 s and thus would have continued into the odor pulse in the muscarine + odor condition. If the odor response was unaffected by muscarine, the muscarine-evoked and odor-evoked increases in GCaMP6f signal should have summed. Instead, the peak calyx ΔF/F during the odor pulse was the same before and after locally applying muscarine, suggesting that the specifically odor-evoked increase in GCaMP6f was decreased by muscarine. Indeed, applying muscarine to the calyx suppressed odor responses in KC axons (**Figure 6F,G**). Given that calyx muscarine suppresses α′β′ axonal odor responses, the decrease in α′β′ KC GCaMP signal in the absence of odor likely reflects suppression of spontaneous action potentials (**Figure 6A,E**), as α′β′ KCs have the highest spontaneous spike rate out of the three subtypes (Groschner et al., 2018; Turner et al., 2008). The increase in calyx Ca^2+^ induced by muscarine alone (without odor) might reflect Ca^2+^ release from internal stores triggered by G_q_ signaling, which then inhibits KC excitability (thus smaller odor responses). Note that muscarine on the calyx is unlikely to reduce KC odor responses via presynaptic inhibition of PNs, because bath muscarine does not affect PN odor responses in the calyx (**Figure 5C**), although we cannot rule out Ca^2+^-independent inhibition. Interestingly, KC axons in many cases show opposite responses to local muscarine application in the horizontal lobe vs. the calyx (e.g., α′β′ and γ KCs in **Figure 6A**), suggesting that mAChR-A may act via different signaling pathways in KC dendrites and axons.

### mAChR-A knockdown prevents training-induced depression of MBON odor responses

The finding that muscarine can locally inhibit Ca^2+^ influx in KC axons suggests that mAChR-A may play a role in the weakening of KC->MBON synapses that underlies olfactory learning. For example, mAChR-A might be required in KC presynaptic terminals to integrate the US (shock) and CS (odor) to induce synaptic plasticity. If so, increasing the strength of the US might overcome the aversive learning defect in mAChR-A knockdown flies. In contrast, if mAChR-A functions only in dendrites and modulates how KCs process the CS, not the US or CS-US integration, changing the US strength should not affect the learning defect in mAChR-A knockdown flies. We tested this by increasing the electric shock strength from 50 V (as in **Figure 1A**) to 90 V (**Figure 7A**). Indeed, increased electric shock strength improved learning (compare scores in **Figure 7A** with **Figure 1A**). In particular, mAChR-A knockdown flies no longer show any learning defect. As only the US input to KC presynaptic terminals was modified while the CS (i.e. odor input to the calyx) was unchanged, this result is consistent with the hypothesis that mAChR-A exerts its effect on learning and memory at KC presynaptic terminals. This result also backs up the imaging results in **Figure 4** showing that mAChR-A knockdown does not affect KC odor coding (i.e., the sparseness or inter-odor correlation of KC population responses).

**Figure 7:**
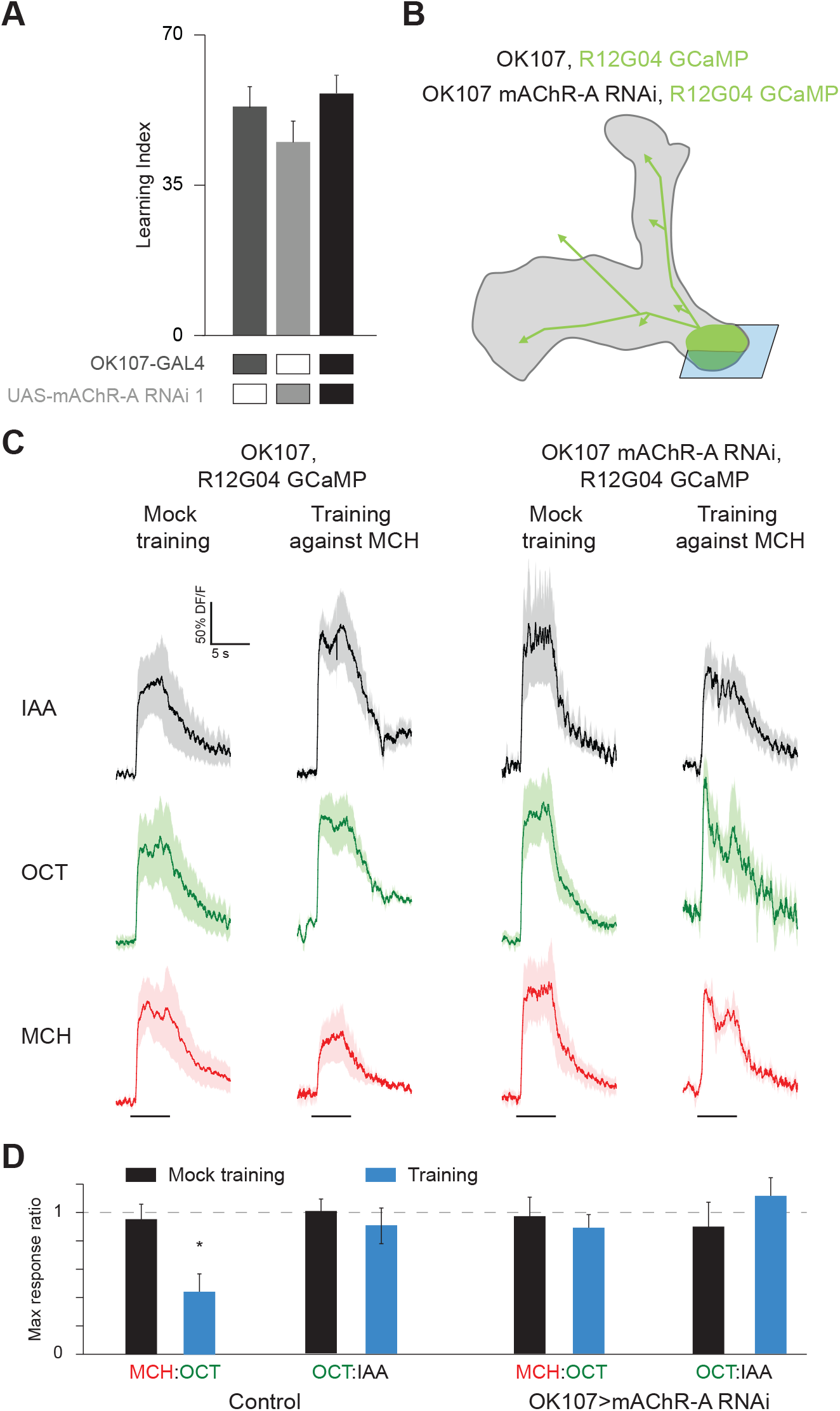
mAChR-A KD prevents aversive conditioning from decreasing the response to the trained odor in MB-MVP2. **(A)** When flies were conditioned against MCH using 90 V electric shock (see Methods), driving mAChR-A RNAi in KCs using OK107-GAL4 did not affect learning compared to controls (mean ± SEM, n (left to right): 52, 49, 51, p > 0.13, Kruskal-Wallis test). For detailed statistical analysis see Table S1. **(B)** Diagram of genotype: mAChR-A RNAi 1 was expressed in KCs with OK107 (gray), while GCaMP6f was expressed in MB-MVP2 with R12G04-LexA (green). The imaging plane is shown in blue. **(C)** Odor responses in MB-MVP2 to isoamyl acetate (IAA, not presented during training), OCT (not shocked during training) and MCH (shocked during training), in control (OK107-GAL4, R12G04-LexA>GCaMP6f, mb247-dsRed) and knockdown (OK107-GAL4>mAChR-A-RNAi 1, R12G04-LexA>GCaMP6f, mb247-dsRed) flies, with mock training (no shock) or training against MCH. Traces show mean (solid line) ± SEM (shaded area). **(D)** MCH:OCT or OCT:IAA ratios of peak ΔF/F values from (C). n = 5. * p<0.05, Mann-Whitney test. Power analysis shows that n = 5 would suffice to detect an effect as strong as the difference between training and mock training in the MCH:OCT ratio, with power 0.9.

We sought to further test whether mAChR-A contributes to the synaptic plasticity underlying aversive olfactory learning. In *Drosophila,* olfactory associative memories are stored by weakening the synapses between KCs and output neurons that lead to the “wrong” behavior. For example, aversive memory requires an output neuron downstream of γ KCs, called MBON-γ1pedc>α/β or MB-MVP2. MB-MVP2 leads to approach behavior, and aversive conditioning reduces MB-MVP2’s responses to the aversively-trained odor (Hige et al., 2015b; Perisse et al., 2016). We tested whether knocking down mAChR-A would prevent this depression. We knocked down mAChR-A in KCs using OK107-GAL4 and UAS-mAChR-A-RNAi 1, and expressed GCaMP6f in MB-MVP2 using R12G04-LexA and lexAop-GCaMP6f (**Figure 7B**). We trained flies in the behavior apparatus and then imaged MB-MVP2 odor responses (3 h after training to avoid cold-shock-sensitive memory). Because overall response amplitudes were variable across flies, for each fly we measured the ratio of the response to MCH (the trained odor) over the response to OCT (the untrained odor). Consistent with previous published results (Hige et al., 2015b; Perisse et al., 2016), in control flies not expressing mAChR-A RNAi, the MCH/OCT ratio was substantially reduced in trained flies relative to mock-trained flies (**Figure 7C**). This was not because the OCT response increased, because there was no difference between trained and mock-trained flies in the ratio of the response to OCT over the response to isoamyl acetate, a ‘reference’ odor that was absent in the training protocol. In contrast, in flies expressing mAChR-A RNAi in KCs, the MCH/OCT ratio was the same between trained and mock-trained flies (**Figure 7C**), indicating that the mAChR-A knockdown impaired the learning-related depression of the KC to MB-MVP2 synapse. These results support the notion that mAChR-A plays a role at the presynaptic terminal of KCs where LTD occurs.

## Discussion

Here we show that mAChR-A is required in γ KCs for aversive olfactory learning and short-term memory in adult *Drosophila*. Knocking down mAChR-A increases KC odor responses, while the mAChR-A agonist muscarine suppresses KC activity at both KC dendrites and axons. Knocking down mAChR-A prevents aversive learning from reducing responses of the MB output neuron MB-MVP2 to the conditioned odor, suggesting that mAChR-A is required for the learning-related depression of KC->MBON synapses.

Why is mAChR-A only required for learning in γ KCs, not αβ or α′β′ KCs? Although our mAChR-A MiMIC gene trap agrees with single-cell transcriptome analysis that α′β′ KCs express less mAChR-A than do γ and αβ KCs (Croset et al., 2018; Davie et al., 2018), transcriptome analysis indicates that α′β′ KCs do express some mAChR-A (**Figure S2**). Moreover, γ and αβ KCs express similar levels of mAChR-A. It may be that the RNAi knockdown is less efficient at affecting the physiology of αβ and α′β′ KCs than γ KCs, whether because the knockdown is less efficient at reducing protein levels, or because αβ and α′β′ KCs have different intrinsic properties or a different function of mAChR-A such that 30% of normal mAChR-A levels is sufficient in αβ and α′β′ KCs but not γ KCs. This interpretation is supported by our finding that mAChR-A RNAi knockdown significantly increases odor responses only in the γ lobe, not the αβ or α′β′ lobes. Alternatively, γ, αβ and α′β′ KCs are thought to be important mainly for short-term memory, long-term memory, and memory consolidation, respectively (Guven-Ozkan and Davis, 2014; Krashes et al., 2007); as we only tested short-term memory, mAChRA may carry out the same function in all KCs, but only its role in γ KCs is required for short-term (as opposed to long-term) memory. Indeed, the key plasticity gene DopR1 is required in γ, not αβ or α′β′ KCs, for short-term memory (Qin et al., 2012).

mAChR-A seems to inhibit KC odor responses, because knocking down mAChRA increases odor responses in the calyx and γ lobe, while activating mAChR-A with bath or local application of muscarine decreases KC odor responses. Some details differ between the genetic and pharmacological results. In particular, while mAChR-A knockdown mainly affects γ KCs, with other subtypes inconsistently affected, bath application of muscarine reduces responses in all KC subtypes, and local axonal application of muscarine reduces responses in the γ and β lobe (for MCH) or all lobes (for OCT). What explains these differences? mAChR-A might be weakly activated in physiological conditions, in which case gain of function would cause a stronger effect than loss of function. Similarly, pharmacological activation of mAChR-A is likely a more drastic manipulation than a two-thirds reduction of mAChR-A mRNA levels. Although we cannot entirely rule out network effects from muscarine application, the effect of muscarine does not stem from PNs or APL (**Figure 5C,D**) and locally applied muscarine would have little effect on neurons outside the mushroom body. A previous report found that local application of ACh at the horizontal lobe did not affect GCaMP5 signal in KC axons, but this experiment may have missed the small dip in GCaMP6f signal (~ -5-10% ΔF/F in the γ lobe), and could not have measured modulation of odor responses as it was done *ex vivo* (Barnstedt et al., 2016).

How does mAChR-A inhibit odor-evoked Ca^2+^ influx in KCs? Given that mAChRA signals through G_q_ when expressed in CHO cells (Ren et al., 2015), that muscarinic G_q_ signaling normally increases excitability in mammals (Caulfield and Birdsall, 1998), and that pan-neuronal artificial activation of G_q_ signaling in *Drosophila* larvae increases overall excitability (Becnel et al., 2013), it may be surprising that mAChR-A inhibits KCs. However, G_q_ signaling may exert different effects on different neurons in the fly brain, and some examples exist of inhibitory G_q_ signaling by mammalian mAChRs. M_1_/M_3_/M_5_ receptors acting via G_q_ can inhibit voltage-dependent Ca^2+^ channels (Gamper et al., 2004; Kammermeier et al., 2000; Keum et al., 2014; Suh et al., 2010), reduce voltage-gated Na+ currents (Cantrell et al., 1996), or trigger surface transport of KCNQ channels (Jiang et al., 2015), thus increasing inhibitory K+ currents. *Drosophila* mAChRA may inhibit KCs through similar mechanisms.

What function does mAChR-A serve in learning and memory? Our results indicate that mAChR-A knockdown prevents the learning-associated weakening of KC-MBON synapses, in particular for MBON-γ1pedc>α/β, aka MB-MVP2 (**Figure 7**). One potential explanation is that the increased odor-evoked Ca^2+^ influx observed in knockdown flies increases synaptic release, which overrides the learning-associated synaptic depression. However, increased odor-evoked Ca^2+^ influx *per se* is unlikely on its own to straightforwardly explain a learning defect, because other genetic manipulations that increase odor-evoked Ca^2+^ influx in KCs either have no effect on, or even improve, olfactory learning. For example, knocking down GABA synthesis in the inhibitory APL neuron increases odor-evoked Ca^2+^ influx in KCs (Lei et al., 2013; Lin et al., 2014) and improves olfactory learning (Liu and Davis, 2008).

Alternatively, mAChR-A may be involved directly in learning-associated synaptic depression. In mammals, where the principal excitatory neurotransmitter is glutamate (vs. ACh in *Drosophila),* metabotropic glutamate receptors (mGluRs) underlie many forms of LTD (Lüscher and Huber, 2010), most notably at the synapse between parallel fibers and Purkinje cells in the cerebellum, where LTD is thought to underlie motor learning (Jörntell and Hansel, 2006). The mushroom body has many architectural similarities to the cerebellum, and the KC-MBON synapse appears to fulfill an analogous role to the parallel fiber–Purkinje cell synapse (Farris, 2011). Using a metabotropic receptor for the main excitatory neurotransmitter to mediate synaptic depression may be yet another conserved feature of this ‘cerebellum-like’ circuit. Although LTD occurs postsynaptically at the parallel–fiber-Purkinje cell synapse and presynaptically at the KC-MBON synapse, mGluRs can also mediate presynaptic LTD (Pinheiro and Mulle, 2008). In addition, mAChRs are also involved in synaptic plasticity and memory in mammals. M_1_ mAChR knockout mice have impaired synaptic plasticity and altered memory (Anagnostaras et al., 2003), and M_1_ mAChRs together with metabotropic glutamate receptors mediate LTD in the hippocampus (Jo et al., 2010; Kamsler et al., 2010; Volk et al., 2007). It may be that mAChR-A similarly co-operates with dopamine receptors to depress KC-MBON synapses. Note that the impaired synaptic depression in mAChR-A knockdown flies could explain the increased odor responses in KC axons. KC-MBON synapses are modulated by individual flies’ experiences before their physiology can be measured experimentally, a phenomenon that requires the plasticity gene *rutabaga* (Hige et al., 2015a). If such plasticity involves depression of KC-MBON synapses and is impaired in mAChR-A knockdown flies, that could explain why mAChR-A knockdown flies have increased odor-evoked Ca^2+^ influx in KC axons. Regardless, mAChR-A likely also inhibits KCs directly, given that muscarine inhibits Ca^2+^ influx in KCs on a time scale of ~1 s.

What role might mAChR-A play in synaptic plasticity? An intriguing possibility is suggested by an apparent paradox: both mAChR-A and the dopamine receptor Damb signal through G_q_ (Himmelreich et al., 2017), but mAChR-A promotes learning while Damb promotes forgetting (Berry et al., 2012). How can G_q_ mediate apparently opposite effects? Perhaps G_q_ signaling aids both learning and forgetting by generally rendering the synapse more labile. Indeed, although *damb* mutants retain memories for longer than wildtype, their initial learning is slightly impaired (Berry et al., 2012); *damb* mutant larvae are also impaired in aversive olfactory learning (Selcho et al., 2009). Although one study reports that knocking down Gq in KCs did not impair initial memory (Himmelreich et al., 2017), the G_q_ knockdown may not have been strong enough; also, that study shocked flies with 90 V shocks, which also gives normal learning in mAChRA knockdown flies (**Figure 7A**).

For mAChR-A to contribute to KC-MBON synaptic depression, which is thought to occur in KC presynaptic terminals, mAChR-A must act at least in part at KC presynaptic terminals. An axonal site of action is supported by our finding that local muscarine application on the horizontal lobe decreases GCaMP6f signal and odor responses in KC axons, including those of γ KCs, the subtype of KCs where mAChR-A is required for learning. Mammalian mAChRs often mediate presynaptic inhibition, most commonly by M_2_-type mAChRs (Allen and Brown, 1993; Bellingham and Berger, 1996; Slutsky et al., 1999) but also in some cases by M_i_-type mAChRs (de Vin et al., 2015; Kamsler et al., 2010; Sheridan and Sutor, 1990). *Drosophila* mAChR-A may combine the signaling pathway of M_1_-type mAChRs (G_q_ rather than G_i/o_) with inhibitory action more typical of M_2_-type mAChRs.

What is the source of ACh which activates mAChR-A and modulates odor responses? In the calyx, cholinergic PNs are certainly a major source of ACh. However, KCs themselves are cholinergic (Barnstedt et al., 2016) and release neurotransmitter in both the calyx and lobes (Christiansen et al., 2011). In the lobes, KCs are the only known source of ACh, as cholinergic MBONs do not have presynaptic specializations in the MB and non-KC intrinsic MB neurons like APL and DPM are not thought to be cholinergic (Haynes et al., 2015; Wu et al., 2013). Thus, mAChRs could function as autoreceptors to prevent excess release of an excitatory neurotransmitter, as in mammals, where metabotropic glutamate receptors mediate presynaptic inhibition on glutamatergic neurons to prevent excess glutamate release (Scanziani et al., 1997). However, mAChRs may also mediate lateral interactions between KCs. Numerous KC-KC synapses have been seen by electron microscopy both in *Drosophila* (Takemura et al., 2017) and other insects (Leitch and Laurent, 1996; Schürmann, 2016; Strausfeld and Li, 1999). Thus, mAChR-A may mediate lateral inhibition, in conjunction with the lateral inhibition provided by the GABAergic APL neuron (Lin et al., 2014), or KC-KC signaling may enhance memory by aiding LTD via mAChR-A.

## Methods

### Fly Strains

Fly strains (see below) were raised on cornmeal agar under a 12 h light/12 h dark cycle and studied 1–10 days post-eclosion. Strains were cultivated at 25 °C unless they expressed temperature-sensitive gene products (GAL80^ts^); in these cases the experimental animals and all relevant controls were grown at 23 °C. To de-repress the expression of RNAi with GAL80^ts^, experimental and control animals were incubated at 31 °C for 7 days. Subsequent behavioral experiments were performed at 25 °C.

Experimental animals carried transgenes over Canton-S chromosomes where possible to minimize genetic differences between strains. The following transgenes were used: *UAS-GCaMP6f, lexAop-GCaMP6f* (Barnstedt et al., 2016; Chen et al., 2013), *UAS-mAChR-A RNAi 1* (TRiP.JF02725, Bloomington #27571), *UAS-mAChR-A RNAi 2* (VDRC ID 101407), *UAS-Dcr-2* (Bloomington #24651), *lexAop-GAL80* (Bloomington #32216), *tub-GAL80^ts^* (McGuire et al., 2003), *mb247-dsRed* (Riemensperger et al., 2005), *GH146-GAL4* (Stocker et al., 1997), *OK107-GAL4* (Connolly et al., 1996), *c305a-GAL4* (Krashes et al., 2007), *mb247-GAL4* (Zars, 2000), *R44E04-LexA* (Bloomington #52736), *R45H04-LexA* (Bräcker et al., 2013), *R12G04-LexA* (Bloomington #52448) (Jenett et al., 2012), *elav-GAL4* (Lin and Goodman, 1994), *NP2631-GAL4, GH146-FLP, tub-FRT-GAL80-FRT, UAS-TNT, UAS-mCherry, mb247-LexA* (Lin et al., 2014), *20xUAS-6xGFP* (Shearin et al., 2014), *UAS-mCD8-GFP*.

### Behavioral Analysis

Behavioral experiments were performed in a custom-built, fully automated apparatus (Claridge-Chang et al., 2009; Lin et al., 2014; Parnas et al., 2013). Single flies were housed in clear polycarbonate chambers (length 50 mm, width 5 mm, height 1.3 mm) with printed circuit boards (PCBs) at both floors and ceilings. Solid-state relays (Panasonic AQV253) connected the PCBs to a 50 V source.

Air flow was controlled with mass flow controllers (CMOSens PerformanceLine, Sensirion). A carrier flow (2.7 l/min) was combined with an odor stream (0.3 l/min) obtained by circulating the air flow through vials filled with a liquid odorant. Odors were prepared at 10 fold dilution in mineral oil. Therefore, liquid dilution and mixing carrier and odor stimulus stream resulted in a final 100 fold dilution of odors. Fresh odors were prepared daily.

The 3 liter/min total flow (carrier and odor stimulus) was split between 20 chambers resulting in a flow rate of 0.15 l/min per half chamber. Two identical odor delivery systems delivered odors independently to each half of the chamber. Air or odor streams from the two halves of the chamber converged at a central choice zone. The 20 chambers were stacked in two columns each containing 10 chambers and were backlit by 940 nm LEDs (Vishay TSAL6400). Images were obtained by a MAKO CMOS camera (Allied Vision Technologies) equipped with a Computar M0814-MP2 lens. The apparatus was operated in a temperature-controlled incubator (Panasonic MIR-154) maintained at 25 °C.

A virtual instrument written in LabVIEW 7.1 (National Instruments) extracted fly position data from video images and controlled the delivery of odors and electric shocks. Data were analyzed in MATLAB 2015b (The MathWorks) and Prism 6 (GraphPad).

A fly’s preference was calculated as the percentage of time that it spent on one side of the chamber. Training and odor avoidance protocols were as depicted in **Figure 1**. The naïve avoidance index was calculated as (preference for left side when it contains air) – (preference for left side when it contains odor). During training, MCH was paired with 12 equally spaced 1.25 s electric shocks at 50 V (Tully and Quinn, 1985). The learning index was calculated as (preference for MCH before training) – (preference for MCH after training). Flies were excluded from analysis if they entered the choice zone fewer than 4 times during odor presentation.

### Functional Imaging

Brains were imaged by two-photon laser-scanning microscopy (Ng et al., 2002; Wang et al., 2003). Cuticle and trachea in a window overlying the required area were removed, and the exposed brain was superfused with carbogenated solution (95% O_2_, 5% CO_2_) containing 103 mM NaCl, 3 mM KCl, 5 mM trehalose, 10 mM glucose, 26 mM NaHCO_3_, 1 mM NaH_2_PO_4_, 3 mM CaCl_2_, 4 mM MgCl_2_, 5 mM N-Tris (TES), pH 7.3.

Odors at 10^-1^ dilution were delivered by switching mass-flow controlled carrier and stimulus streams (Sensirion) via software controlled solenoid valves (The Lee Company). Flow rates at the exit port of the odor tube were 0.5 or 0.8 l/min.

Fluorescence was excited by a Ti-Sapphire laser centered at 910 nm, attenuated by a Pockels cell (Conoptics) and coupled to a galvo-resonant scanner. Excitation light was focussed by a 20X, 1.0 NA objective (Olympus XLUMPLFLN20XW), and emitted photons were detected by GaAsP photomultiplier tubes (Hamamatsu Photonics, H10770PA-40SEL), whose currents were amplified and transferred to the imaging computer. Two imaging systems were used, #1 for **Figures 3–6** except 5C, and #2 for **Figure 5C** and **Figure 7**, which differed in the following components: laser (1: Mai Tai eHP DS, 70 fs pulses; 2: Mai Tai HP DS, 100 fs pulses; both from Spectra-Physics); microscope (1: Movable Objective Microscope; 2: DF-Scope installed on an Olympus BX51WI microscope; both from Sutter); amplifier for PMT currents (1: Thorlabs TIA-60; 2: Hamamatsu HC-130-INV); software (1: ScanImage 5; 2: MScan 2.3.01). Volume imaging on System 1 was performed using a piezo objective stage (nPFocus400, nPoint). Muscarine was applied locally by pressure ejection from patch pipettes (resistance ~10 MOhm; capillary inner diameter 0.86 mm, outer diameter 1.5 mm; concentration in pipette 20 mM; pressure 12.5 psi) using a Picospritzer III (Parker). A red dye was added to the pipette to visualize the ejected fluid (SeTau-647, SETA BioMedicals) (Podgorski et al., 2012).

Movies were motion-corrected in X-Y using the moco ImageJ plugin (Dubbs et al., 2016), with pre-processing to collapse volume movies in Z and to smooth the image with a Gaussian filter (standard deviation = 4 pixels; the displacements generated from the smoothed movie were then applied to the original, unsmoothed movie), and motion-corrected in Z by maximizing the pixel-by-pixel correlation between each volume and the average volume across time points. ΔF/F, activity maps, sparseness and inter-odor correlation were calculated as in (Lin et al., 2014). We excluded non-responsive flies and flies whose motion could not be corrected.

### Structural Imaging

Brain dissections, fixation, and immunostaining were performed as described (Pitman et al., 2011; Wu and Luo, 2006). To visualize native GFP fluorescence, dissected brains were fixed in 4% (w/v) paraformaldehyde in PBS (1.86 mM NaH_2_PO_4_, 8.41 mM Na_2_HPO_4_, 175 mM NaCl) and fixed for 20 min at room temperature. Samples were washed for 3×20 min in PBS containing 0.3% (v/v) Triton-X-100 (PBT). The neuropil was counterstained with nc82 (DSHB) and goat anti-mouse Alexa 647 or Alexa 564. Primary antisera were applied for 1-2 days and secondary antisera for 1-2 days in PBT at 4 °C, followed by embedding in Vectashield. Images were collected on a Leica TCS SP5, SP8, or Nikon A1 confocal microscope and processed in ImageJ.

APL expression of tetanus toxin was scored by widefield imaging of mCherry. mCherry expression in APL was distinguished from 3XP3-driven dsRed from the GH146-FLP transgene by using separate filter cubes for dsRed (49004, Chroma: 545/25 excitation; 565 dichroic; 605/70 emission) and mCherry (LED-mCherry-A-000, Semrock: 578/21 excitation; 596 dichroic; 641/75 emission).

### Statistics

Statistical analyses were carried out in GraphPad Prism as described in figure legends and **Table S1**. In general, no statistical methods were used to predetermine sample sizes, but where conclusions were drawn from the absence of a statistically significant difference, a power analysis was carried out in G*Power to confirm that the sample size provided sufficient power to detect an effect of the expected size. The experimenter was blind to which hemispheres had APL neurons expressing tetanus toxin before post-experiment dissection (**Figure 5**) but not otherwise.

## Acknowledgments

We thank Vincent Croset, Christoph Treiber and Scott Waddell for sharing mAChR-A expression data before publication. We thank Oren Schuldiner, Andreas Thum, Scott Waddell, the Bloomington Stock Center, the Vienna *Drosophila* RNAi Center, and the Kyoto *Drosophila* Genetic Resource center for fly strains. We thank Anton Nikolaev for comments on the manuscript. This work was supported by the European Research Council (676844, MP; 639489, AL).

## Author contributions

NB: Methodology, investigation, formal analysis, writing–review & editing, visualization. HA: Methodology, investigation, formal analysis, writing-original draft, writing—review & editing, visualization. AAA: Methodology, investigation, formal analysis, writing–review & editing, visualization. ER: Methodology, investigation, formal analysis, software, writing–review & editing, visualization. HL: Investigation, formal analysis, writing-review & editing. WH: Investigation, writing-review & editing, visualization. ACL: Conceptualization, methodology, investigation, formal analysis, software, writing–original draft, writing–review & editing, visualization, supervision, funding acquisition. MP: Initiated the project, conceptualization, methodology, investigation, formal analysis, software, writing–original draft, writing–review & editing, visualization, supervision, funding acquisition.

---

**Figure S1:**
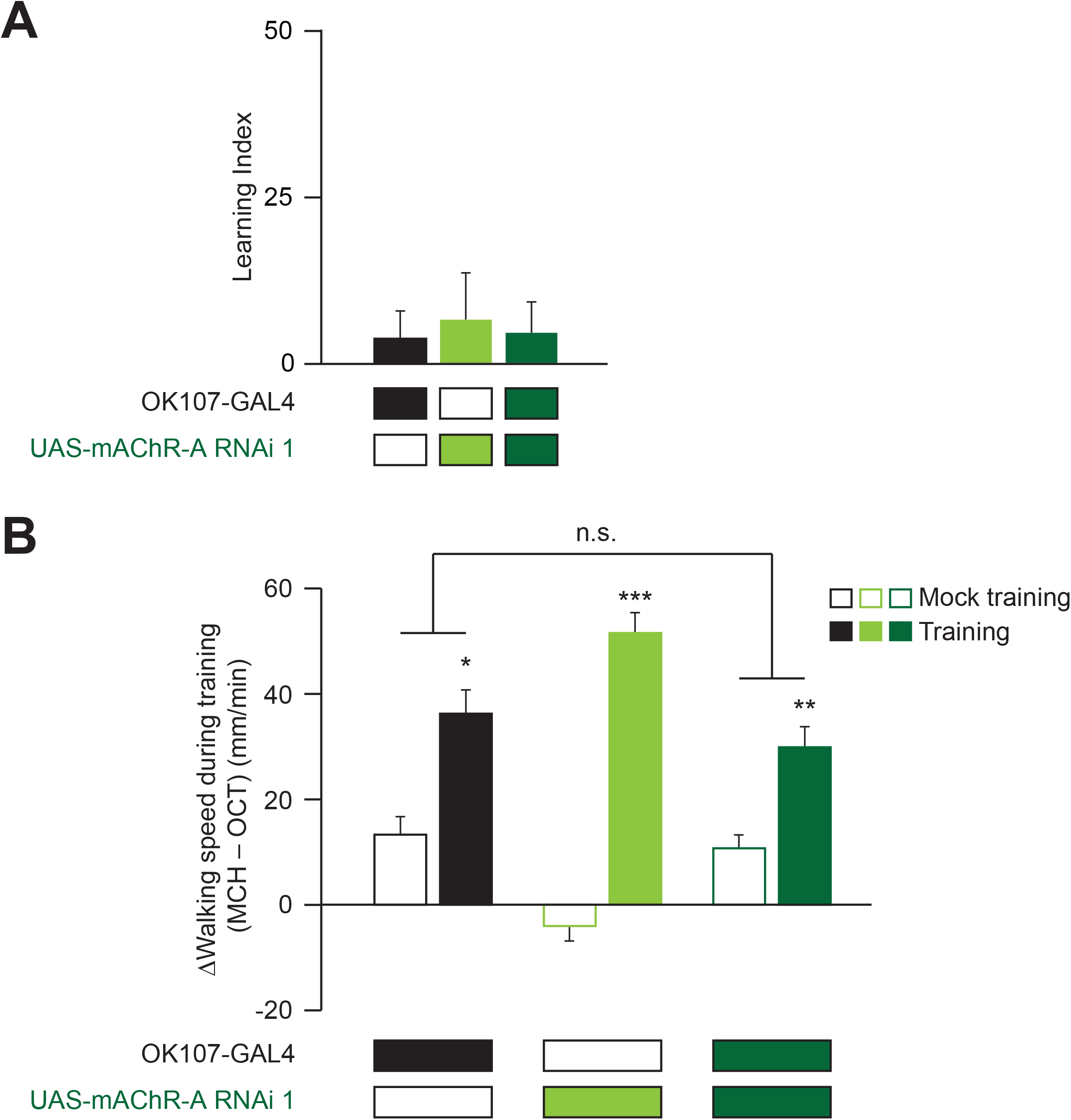
Mock trained flies do not change odor preference over the course of the training protocol. **(A)** Flies were subjected to the same protocol as in Figure 1 but without electric shock. As expected the flies do not change their odor preference and have a learning index which is not statistically different from 0 (n (left to right): 79, 73, 71; p > 0.3, one-sample t-test). For detailed statistical analysis see Table S1. **(B)** Sensitivity to shock (extent to which flies walk faster while being shocked) is not affected by knocking down mAChR-A in KCs. Shown here is walking speed during training (time = 5-6 and 7-8 min in Figure 1B), taking the difference between speed during MCH (CS+) and speed during OCT (CS–). In mock training, the difference is close to zero, but during training, when MCH is paired with shock, flies walk much faster in MCH (* p < 0.05, ** p < 0.01, *** p < 0.001, Mann-Whitney test with Bonferroni correction, comparing training vs. mock training). The effect of shock is not significantly different between OK107 alone and OK107>mAChR-A-RNAi flies (n.s.: p = 0.44 for interaction between genotype and training vs. mock training, 2-way ANOVA). n (left to right): 79, 99, 79, 79, 139, 159.

**Figure S2:**
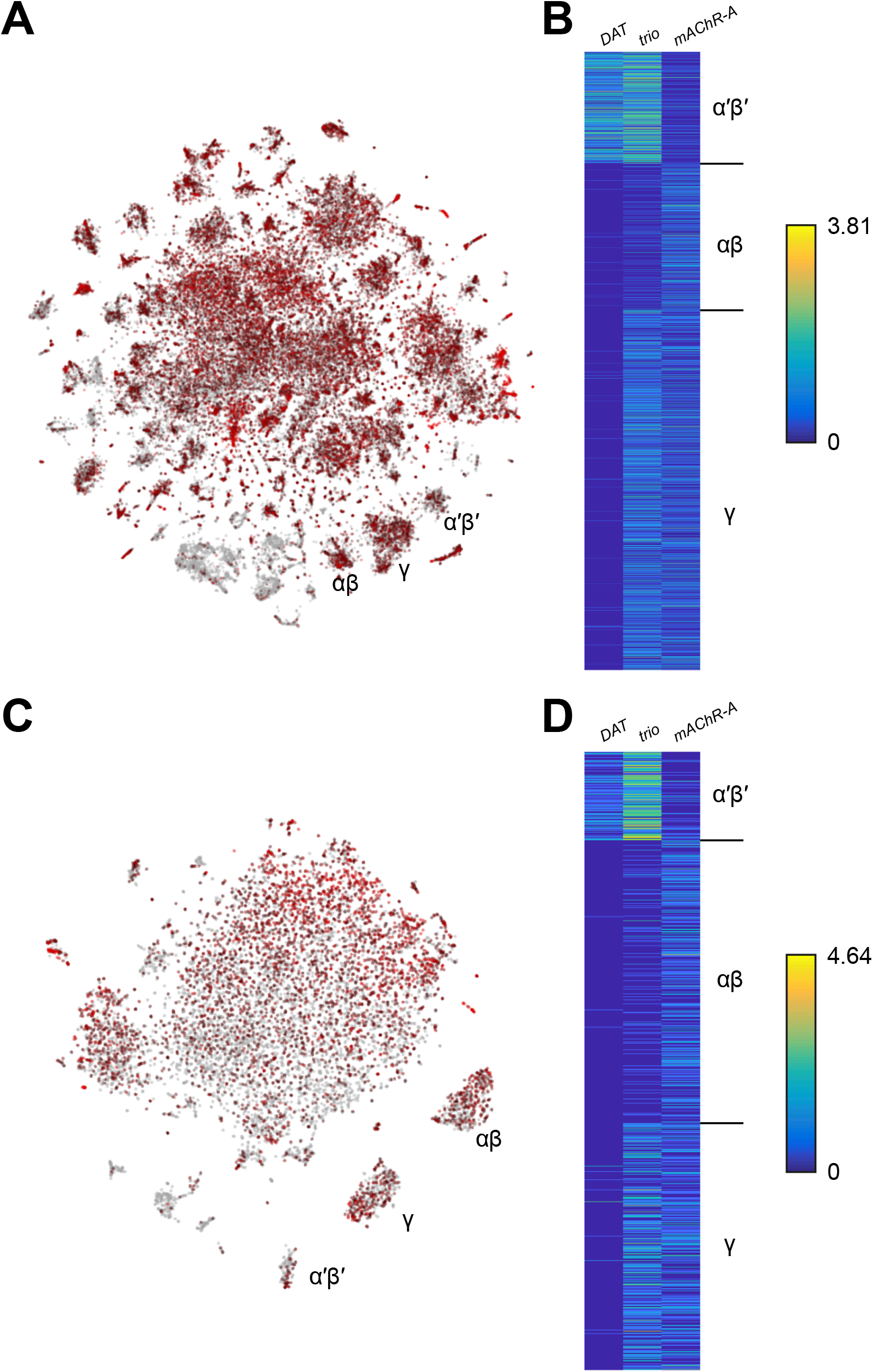
Expression of mAChR-A from single-cell transcriptome profiling. **(A)** Data from Davie et al., 2018. 56,902 *Drosophila* brain cells arranged according to their single-cell transcriptome profiles, along the top 2 principal components using t-SNE. Red coloring indicates expression of mAChR-A. KC subtype clusters are labeled as identified in Davie et al., 2018. **(B)** Expression of *DAT* (marker for α′β′ KCs), *trio* (marker for α′β′ and γ KCs), and *mAChR-A* for cells identified as α′β′, αβ and γ KCs in Davie et al., 2018. mAChR-A expression is higher in αβ and γ KCs compared to α′β′ KCs. **(C)** As in A but with data from Croset et al., 2018 (10,286 *Drosophila* brain cells). **(D)** As in B but with data from Croset et al., 2018. Images screenshotted and raw data downloaded from SCope (http://scope.aertslab.org) on 24 June 2018.

**Figure S3:**
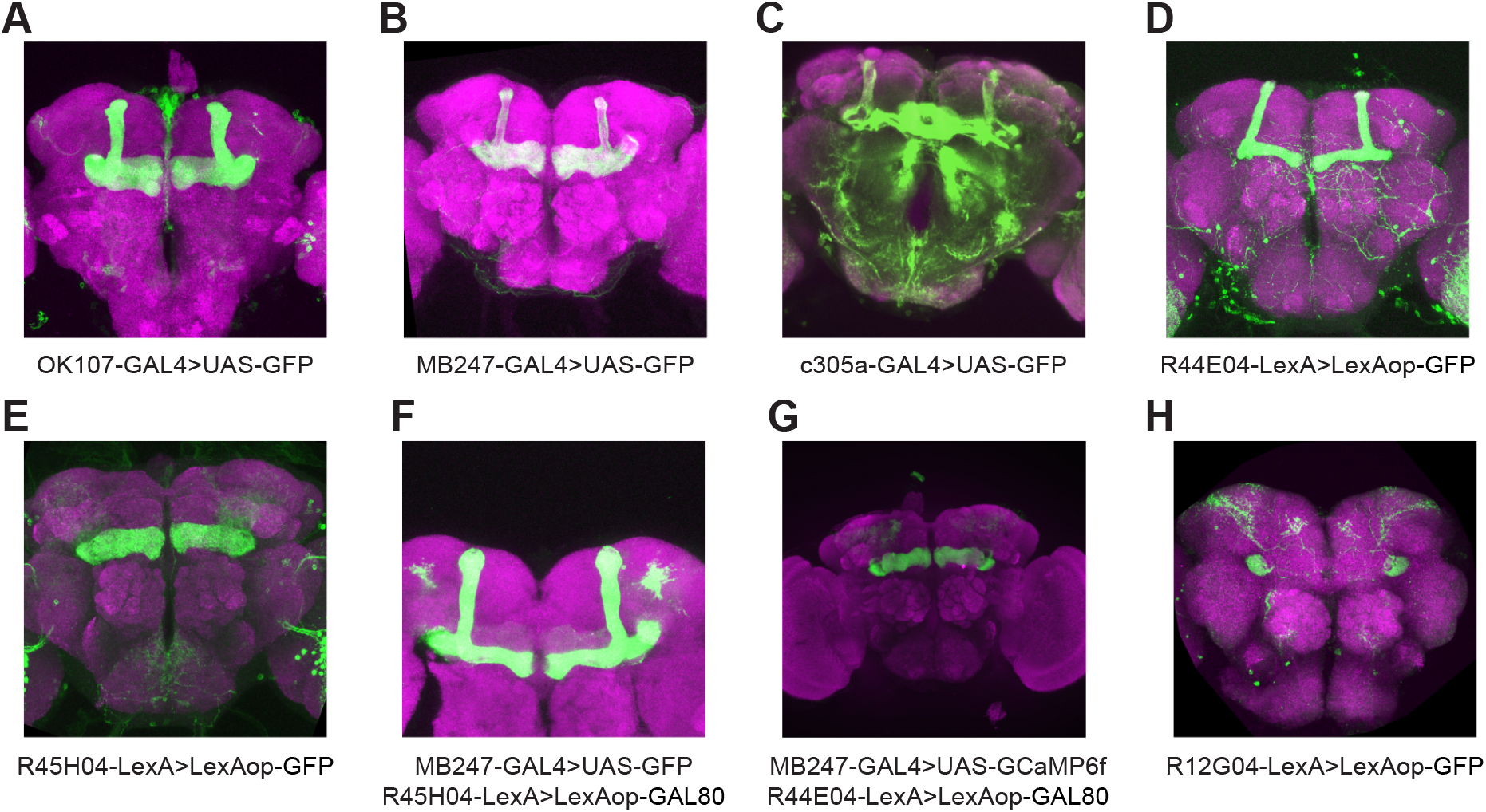
Expression patterns of GAL4 and LexA driver lines used in this study. GFP expression was driven by the named GAL4 or LexA driver lines and the general neuropil was stained with an antibody to NC82 (magenta). Images are maximum-intensity Z-projections of confocal stacks. Panels A-D, G show only the planes of the mushroom body lobes and peduncle to more clearly show which lobes are labeled. **(A)** OK107-GAL4 labels all KCs. **(B)** MB247-GAL4 labels αβ and γ KCs. **(C)** c305a-GAL4 labels α′β′ KCs. **(D)** R44E04-LexA labels αβ KCs. **(E)** R45H04-LexA strongly labels γ KCs. **(F)** Silencing MB247-GAL4 expression in γ KCs by using R45H04-LexA to drive lexAop-GAL80 in γ KCs results in fairly specific expression in αβ KCs. **(G)** Silencing MB247-GAL4 expression in αβ KCs by using R44E04 to drive lexAop-GAL80 in αβ KCs results in fairly specific expression in γ KCs. **(H)** R12G04-GAL4 labels MBON-γ1pedc>α/β, aka MB-MVP2.

**Figure S4:**
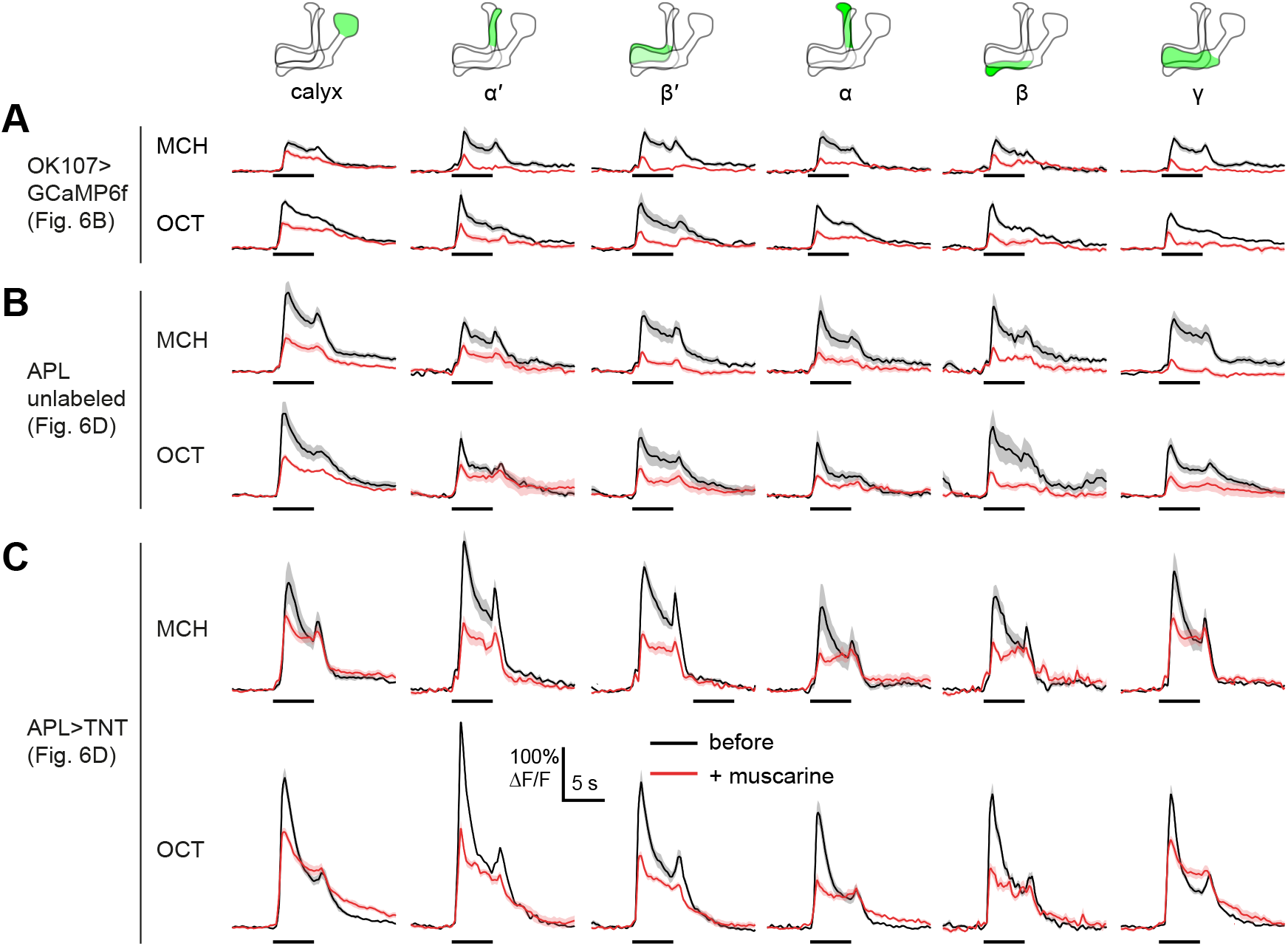
Odor responses of all lobes are reduced by bath muscarine. Extended data for **Figure 5**. Odor responses in OK107-GAL4>GCaMP6f flies **(A)**, control APL unlabeled hemispheres **(B)**, and APL>TNT hemispheres **(C)**, before (black) and after (red) adding 10 μM muscarine in the bath. Data are mean (solid line) ± SEM (shaded area); diagrams illustrate which region of the MB was analyzed; horizontal lines indicate the odor pulse. These are the traces for the summary data shown in **Figure 5B,D**.

**Figure S5:**
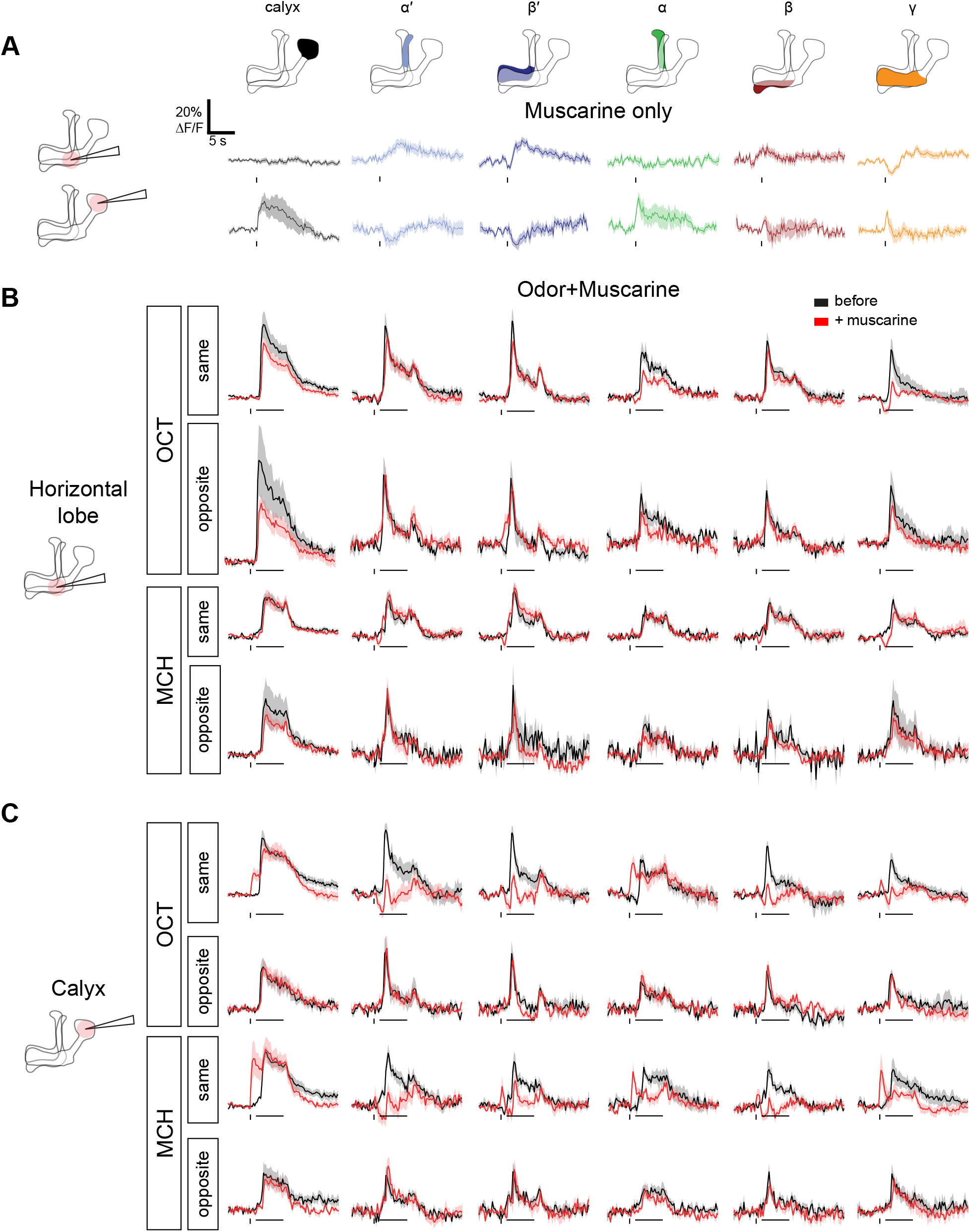
Muscarine differentially affects the MB regions when it is locally applied to the calyx or horizontal lobe. Average ΔF/F GCaMP6f traces of the different MB regions of OK107>GCaMP6f flies that only received the muscarine pulse **(A)** or received an odor pulse (MCH or OCT) before (black) or after (red) 10 ms pulse of 20 mM muscarine **(B,C)**. Panel **A** is duplicated from **Figure 6A**; panels **B,C** are the traces corresponding to the **Figure 6G**. Muscarine was applied either in the horizontal lobe or calyx, 1 s before the odor pulse where applicable. Traces are from the same side or the opposite side that muscarine was applied. Data are mean (solid line) ± SEM (shaded area). Horizontal bars indicate odor pulse timing and duration. Vertical bars indicate timing of muscarine pulse. n, by number of hemispheres (number of flies): Horizontal lobe same side MCH 13 (8), OCT 12 (8), opposite side MCH 6 (3), OCT 6 (3), muscarine alone 15 (8). Calyx same side MCH 6 (4), OCT 7 (5), opposite side MCH 5 (3), OCT 5 (3), muscarine alone 7 (5). n for odor + muscarine traces differs between this figure vs. **Figure 6G** because a scripting error meant that for 10 of the recordings, the odor pulse paired with muscarine presentation lasted 4 s, not 5 s. These data are included in **Figure 6G**, because the peak ΔF/F always occurred within the first 4 s of the odor pulse and thus was unaffected by the scripting error, but they are excluded from traces shown in this figure because the stimuli do not match.

**Table S1.**
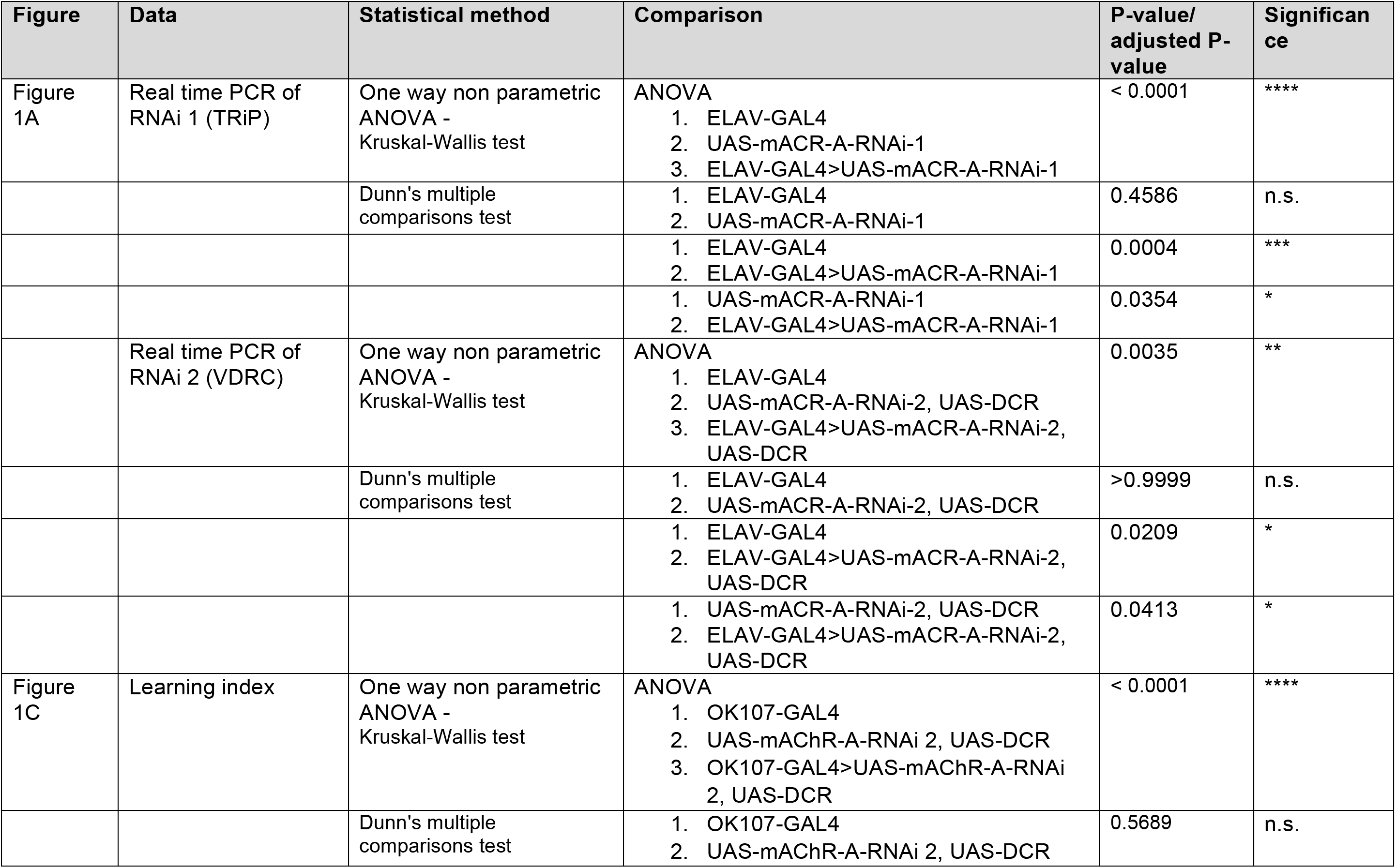
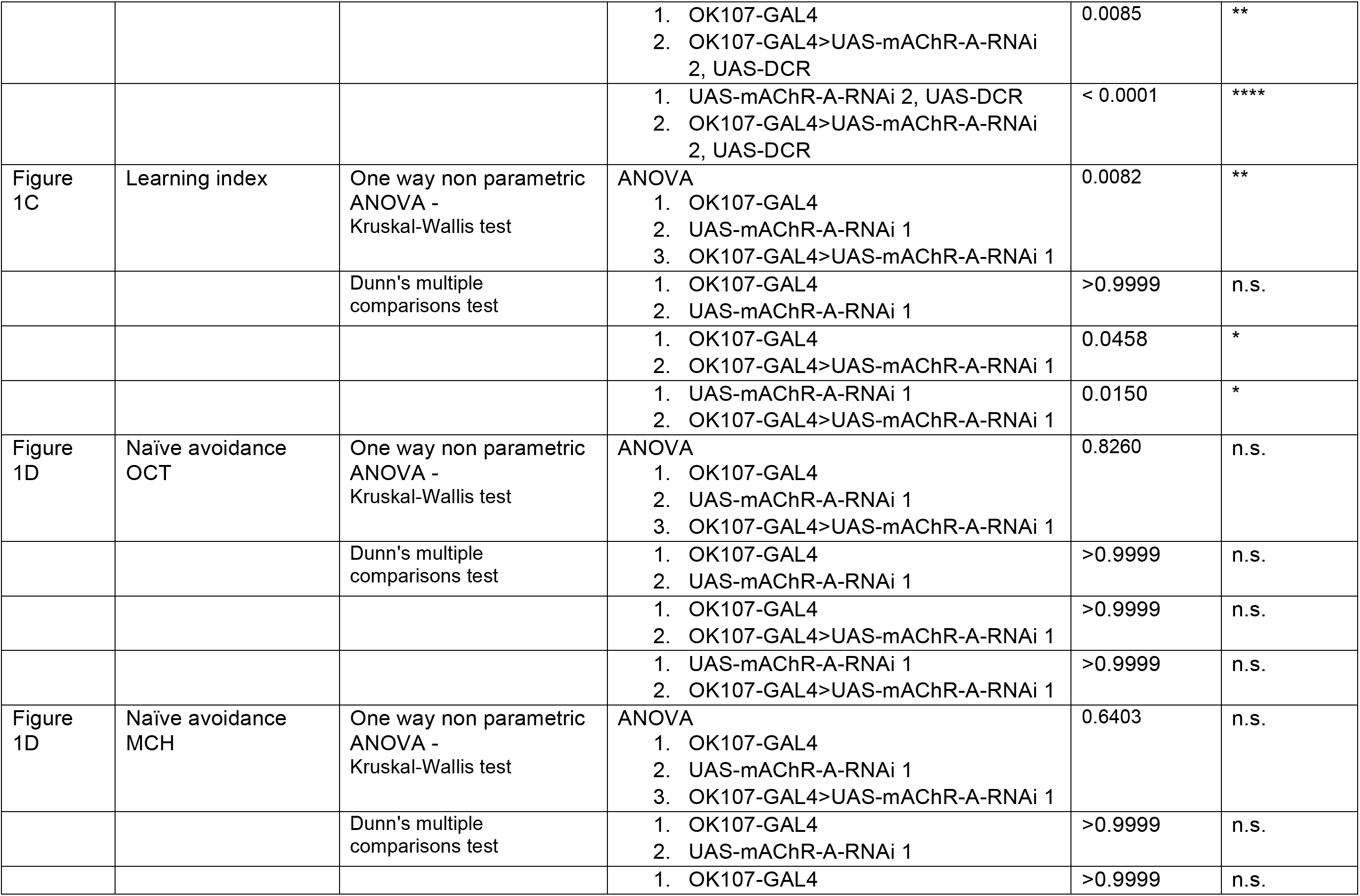
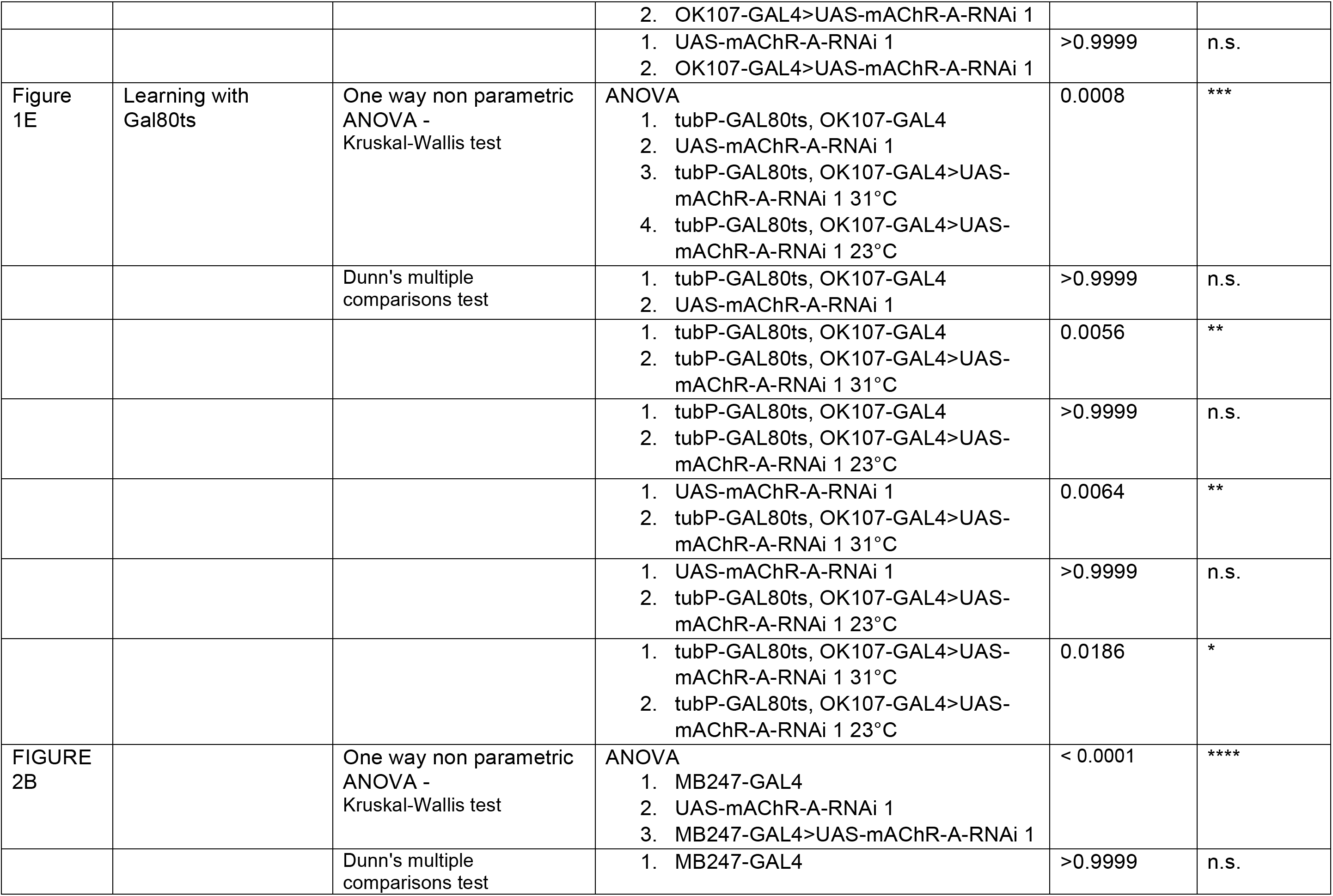
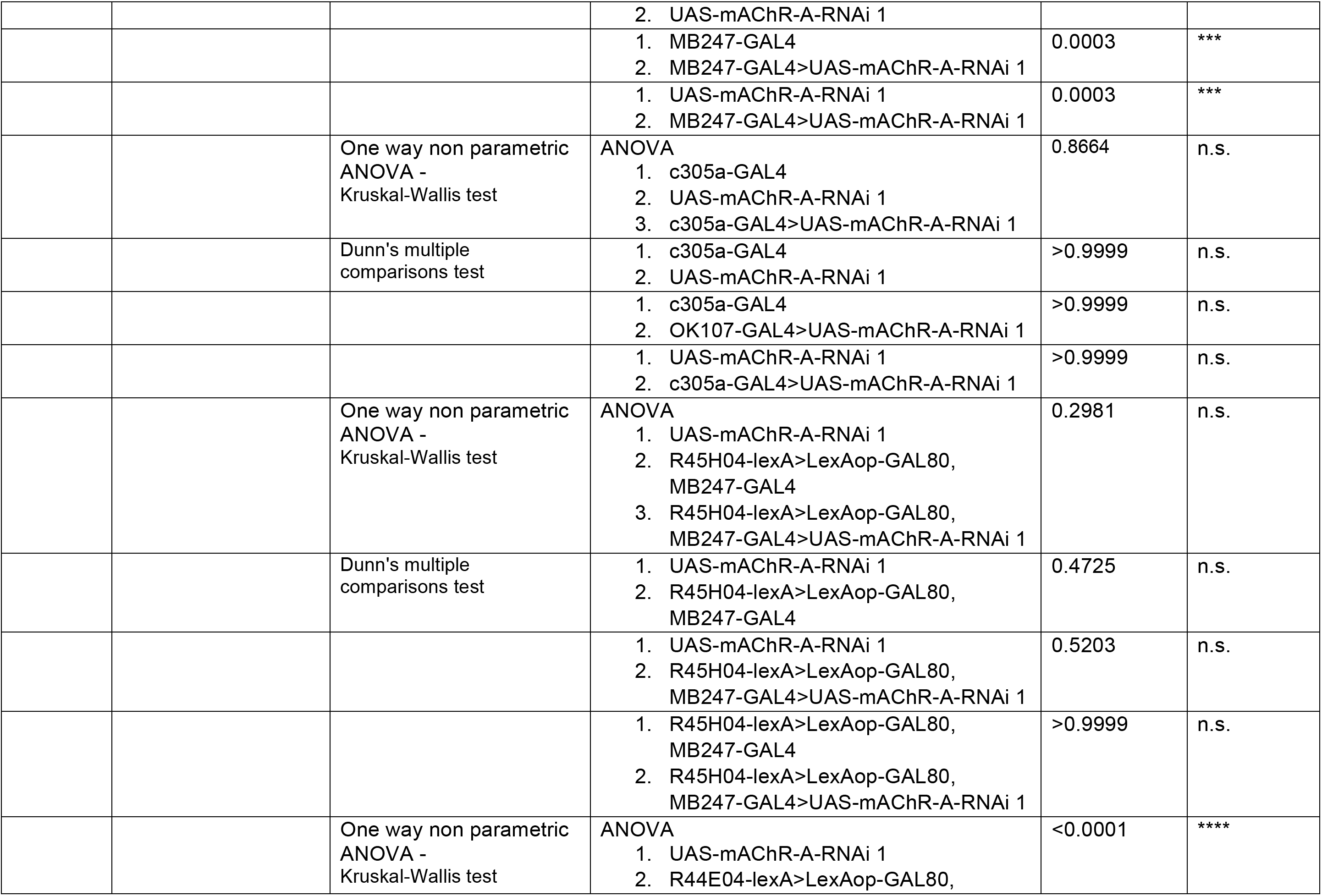
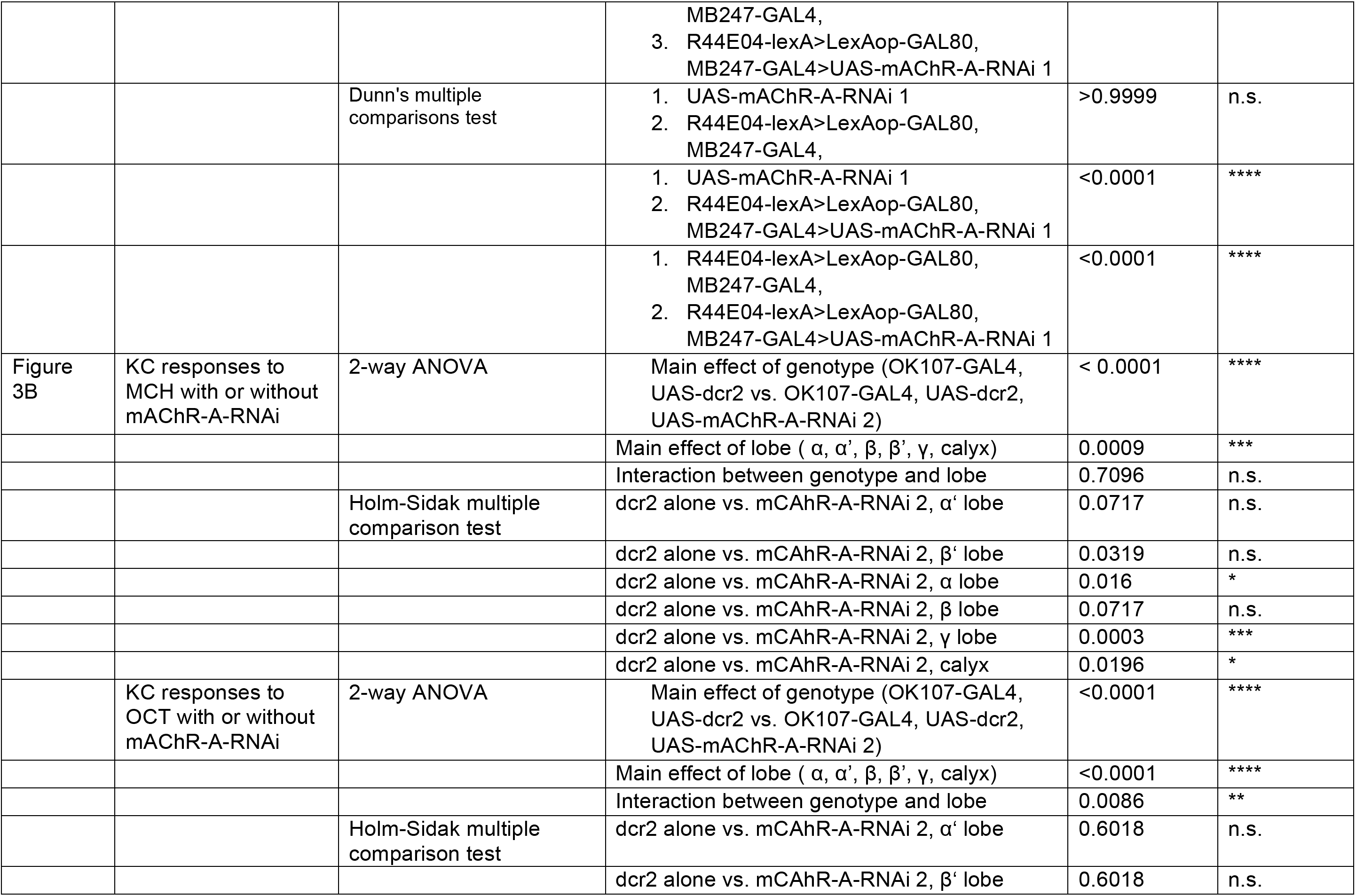
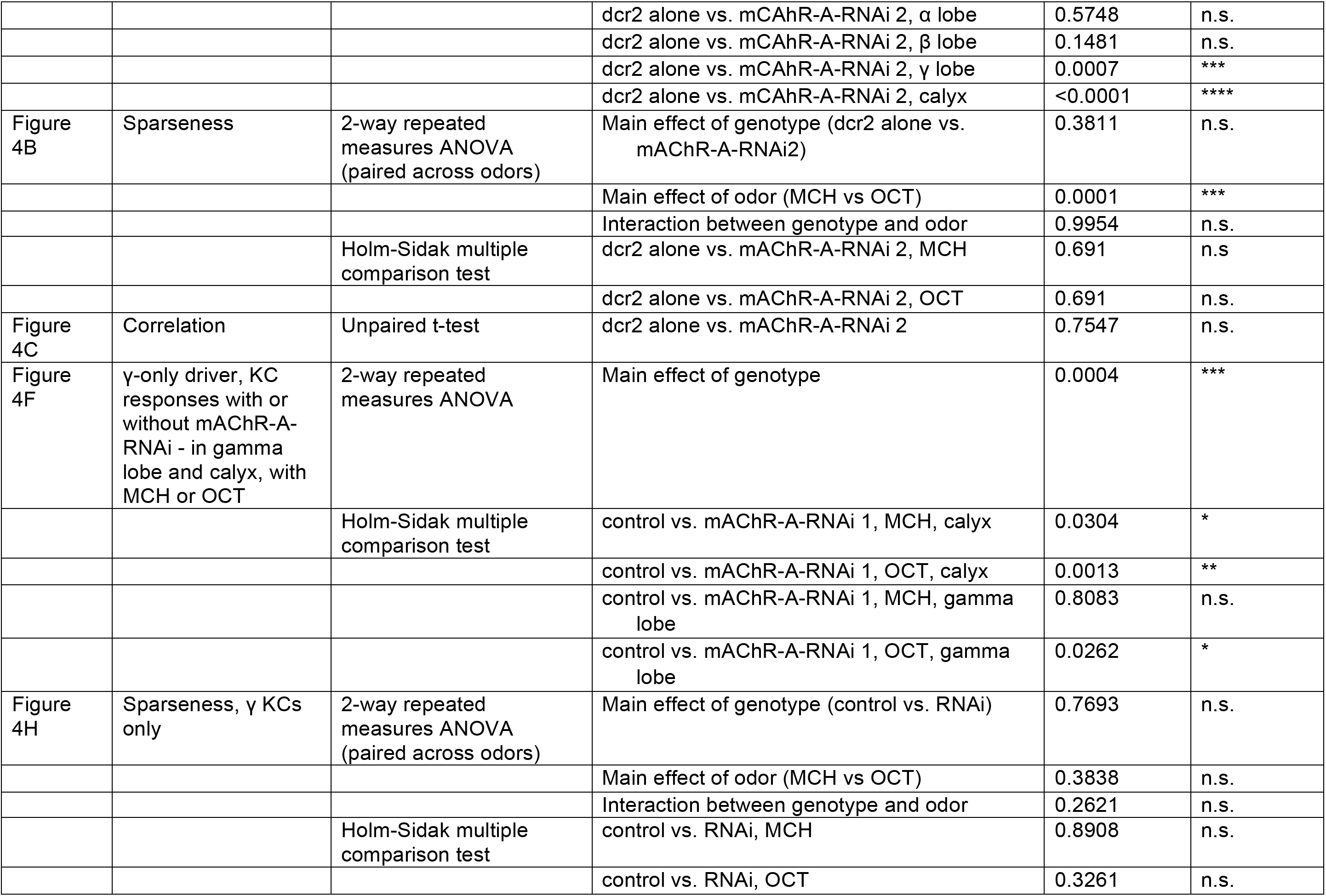
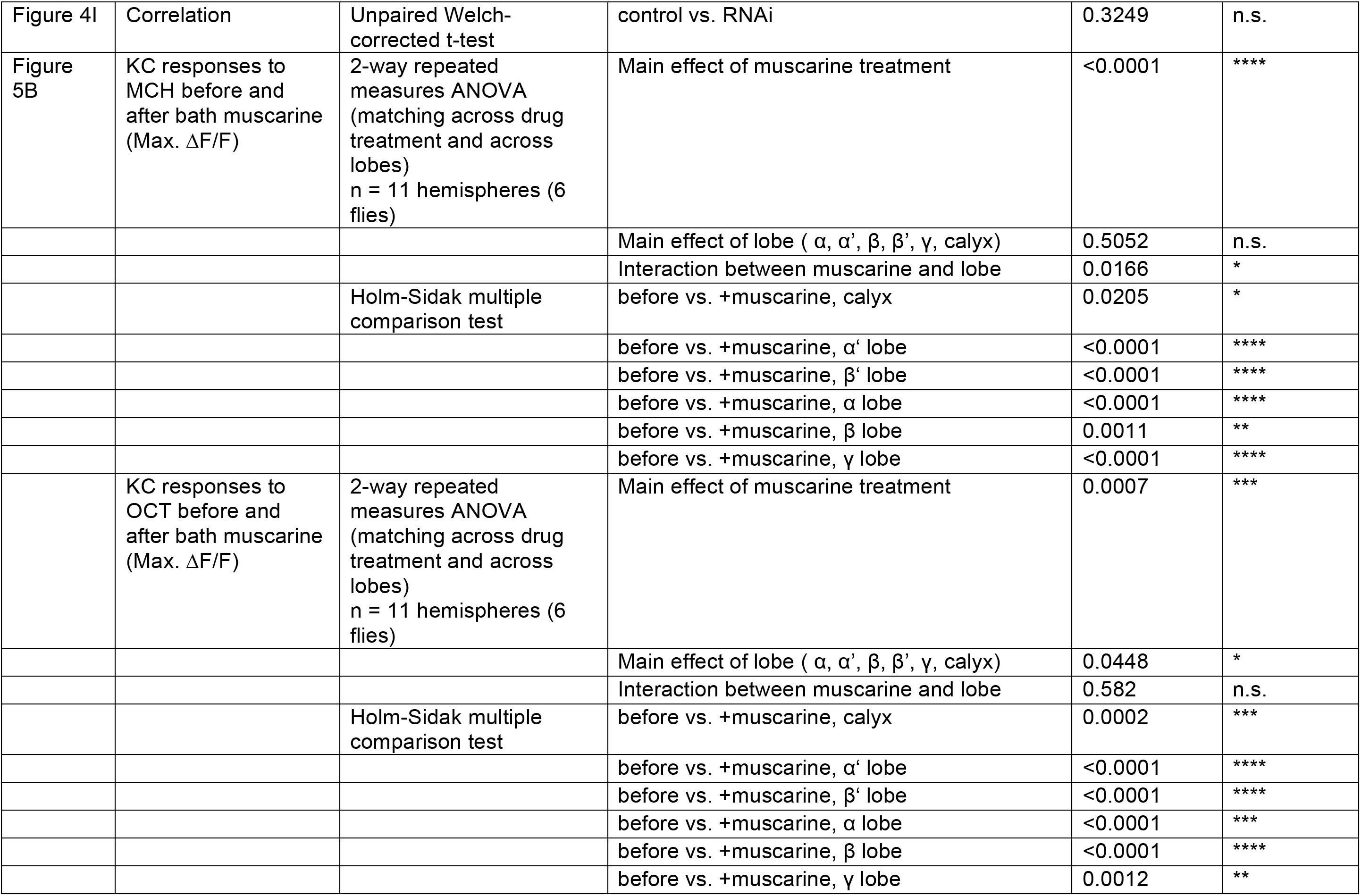
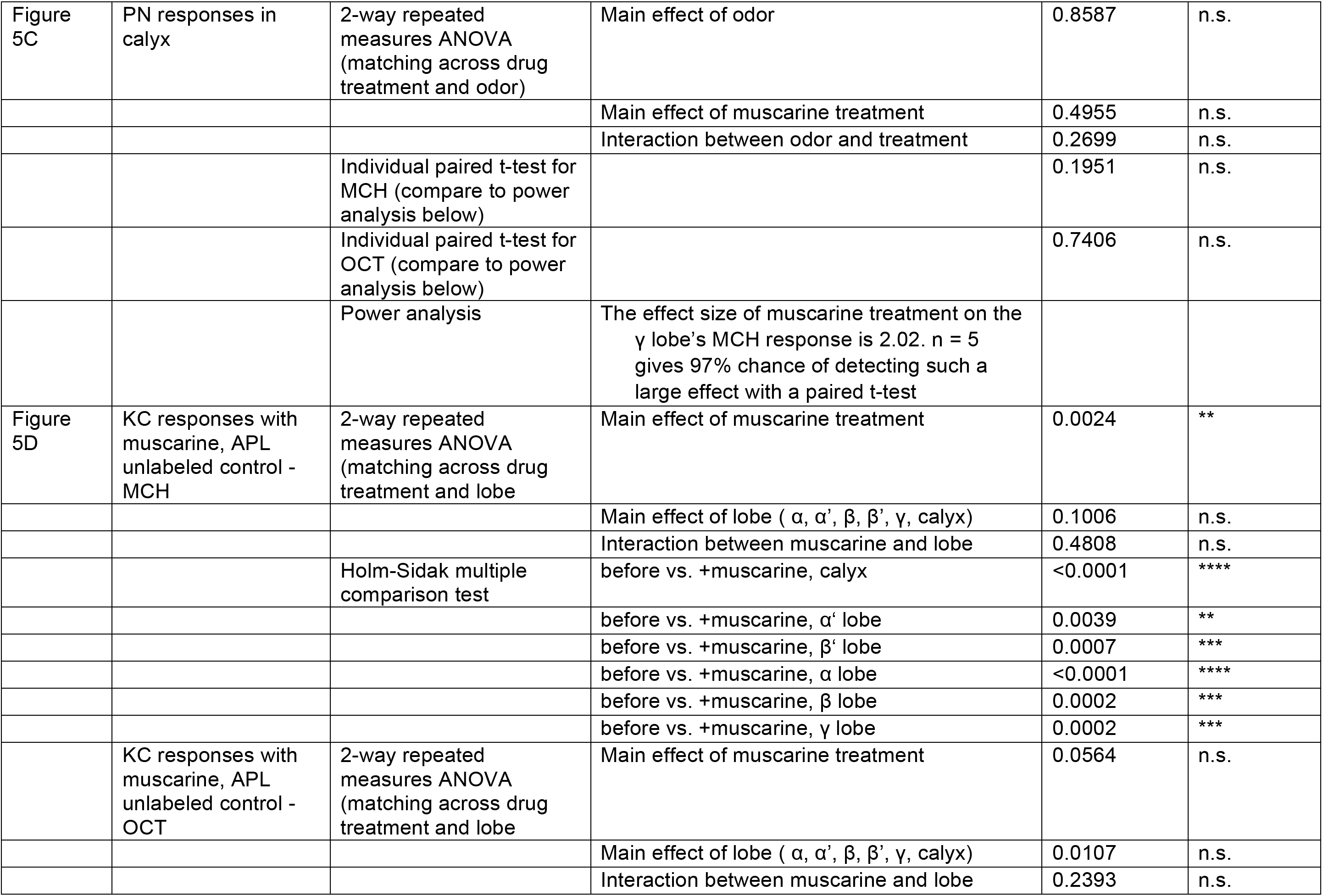
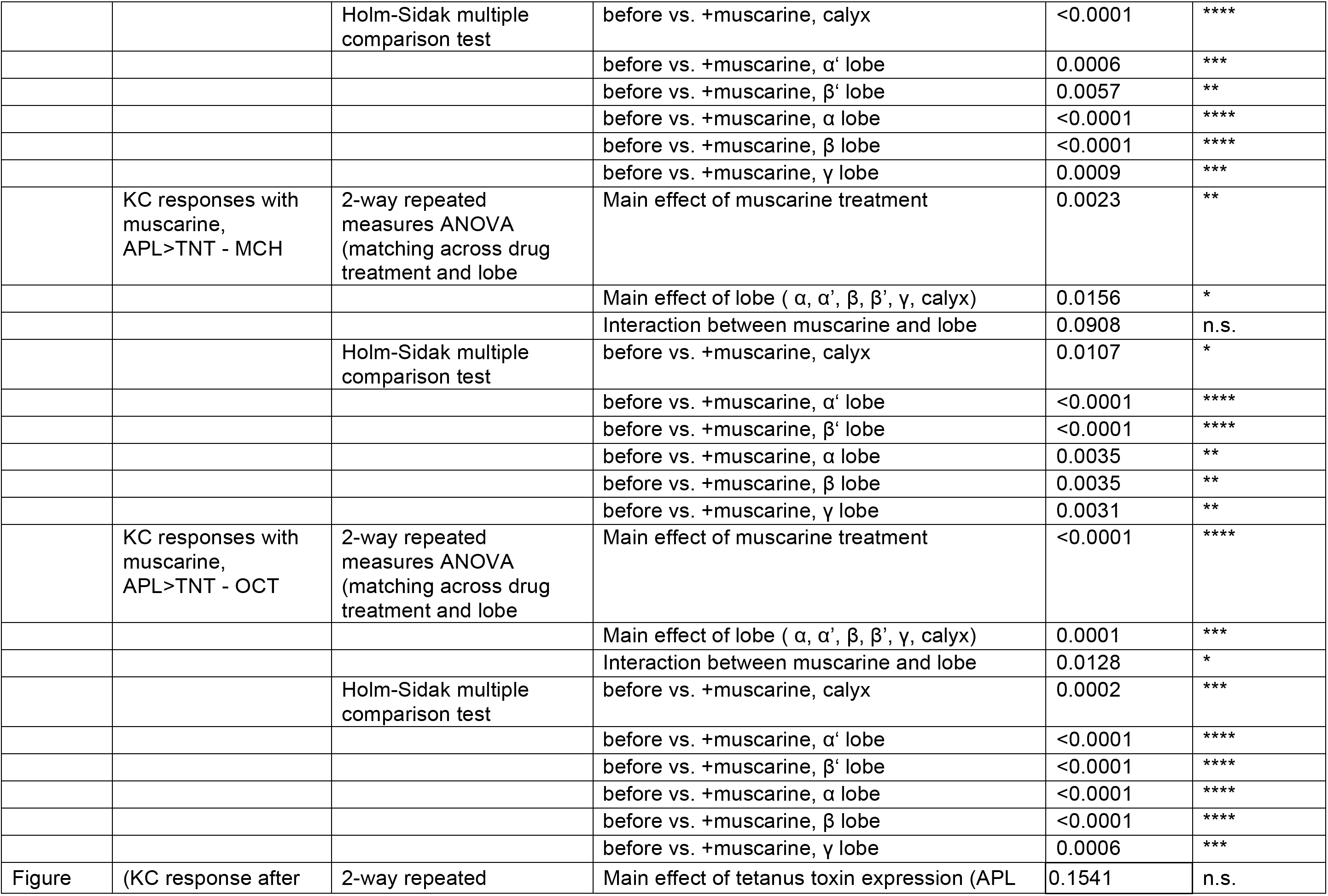
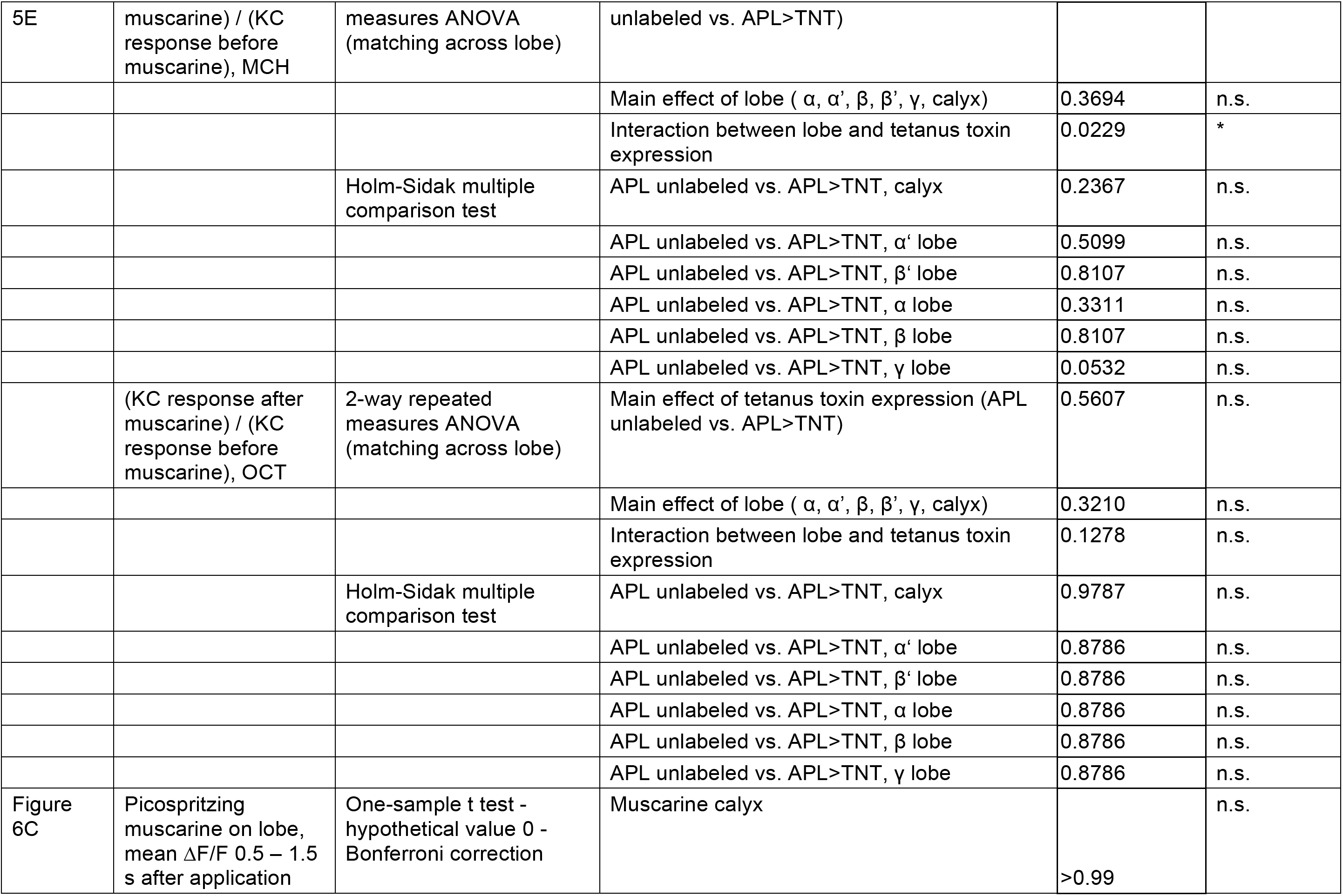
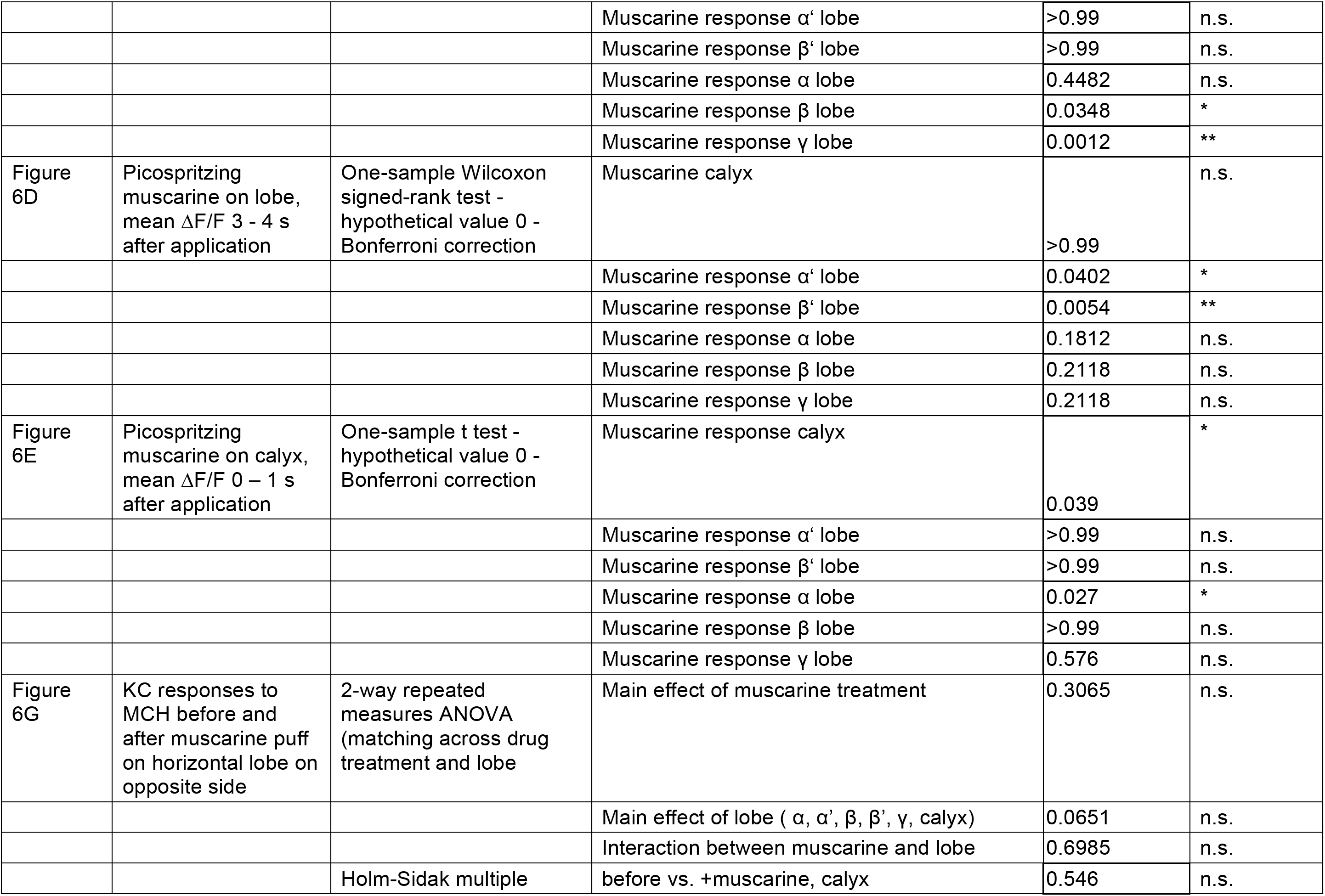
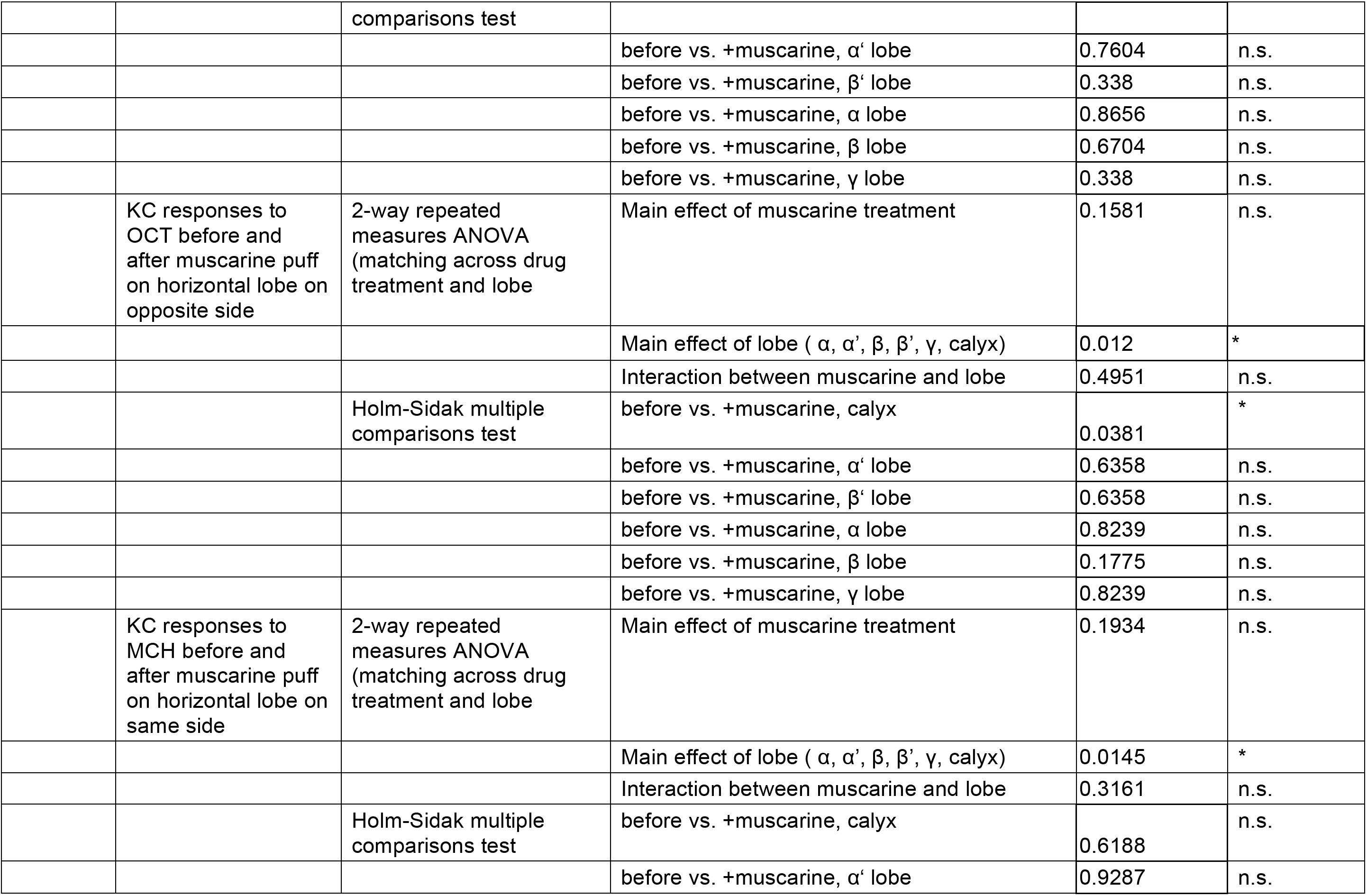
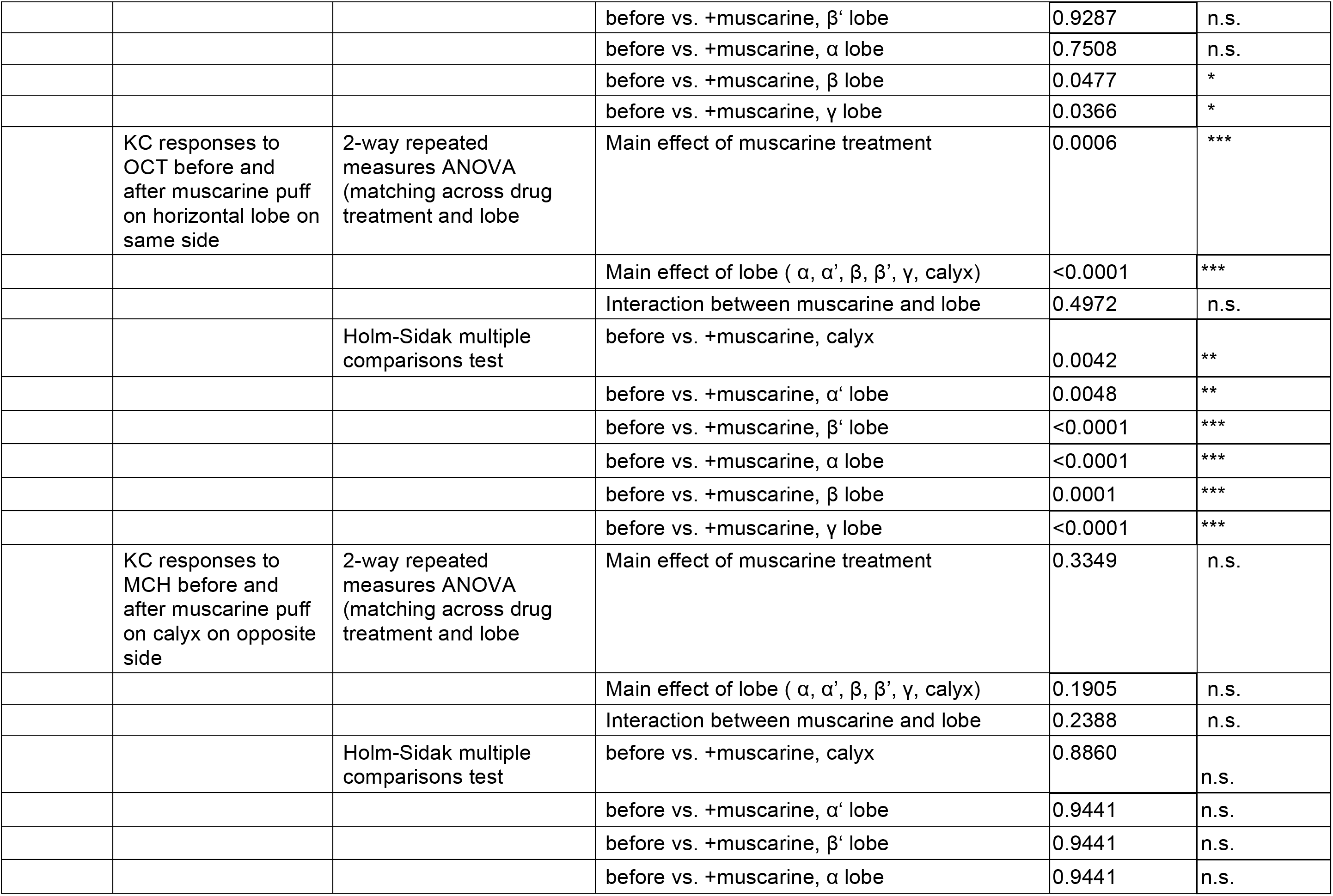
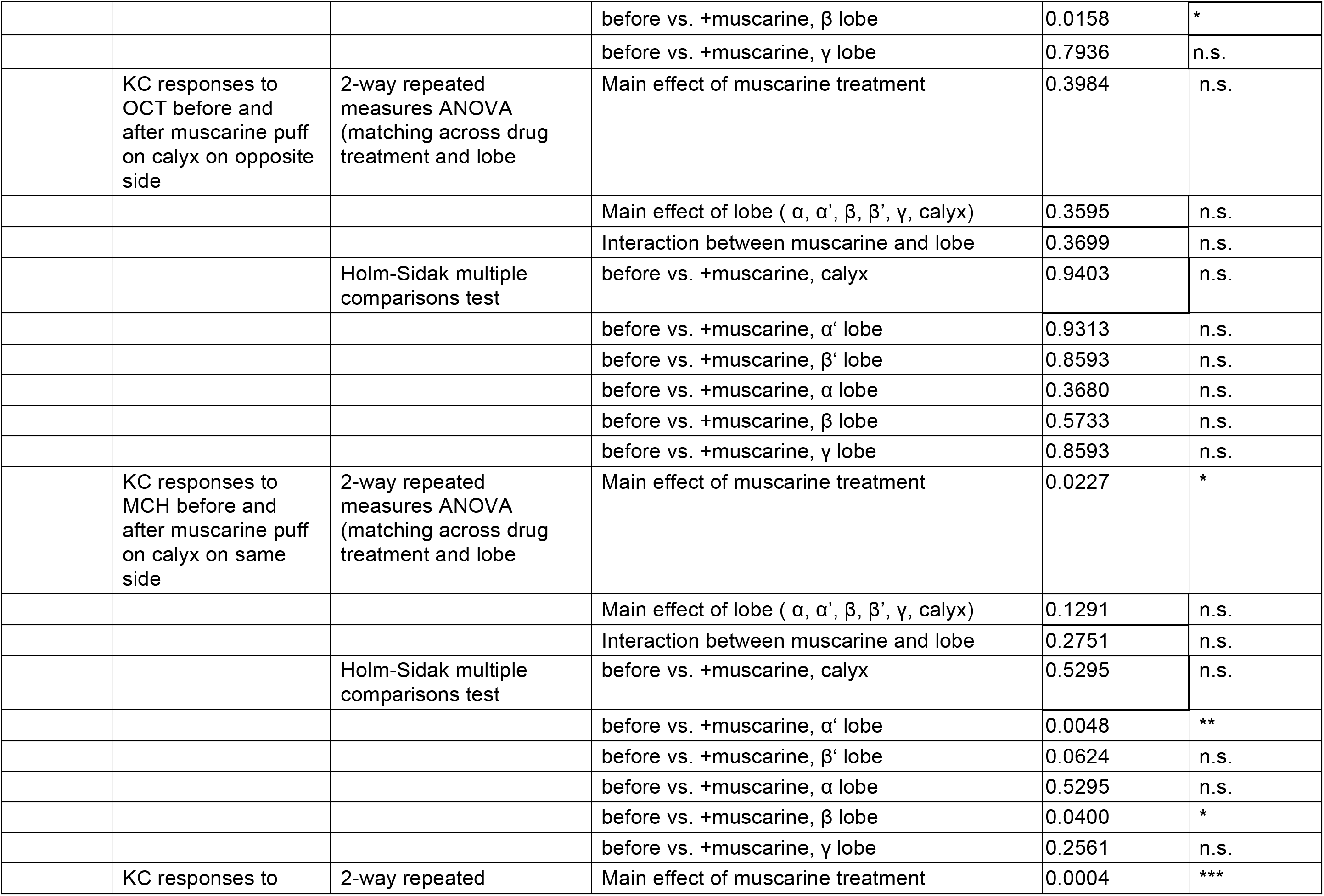
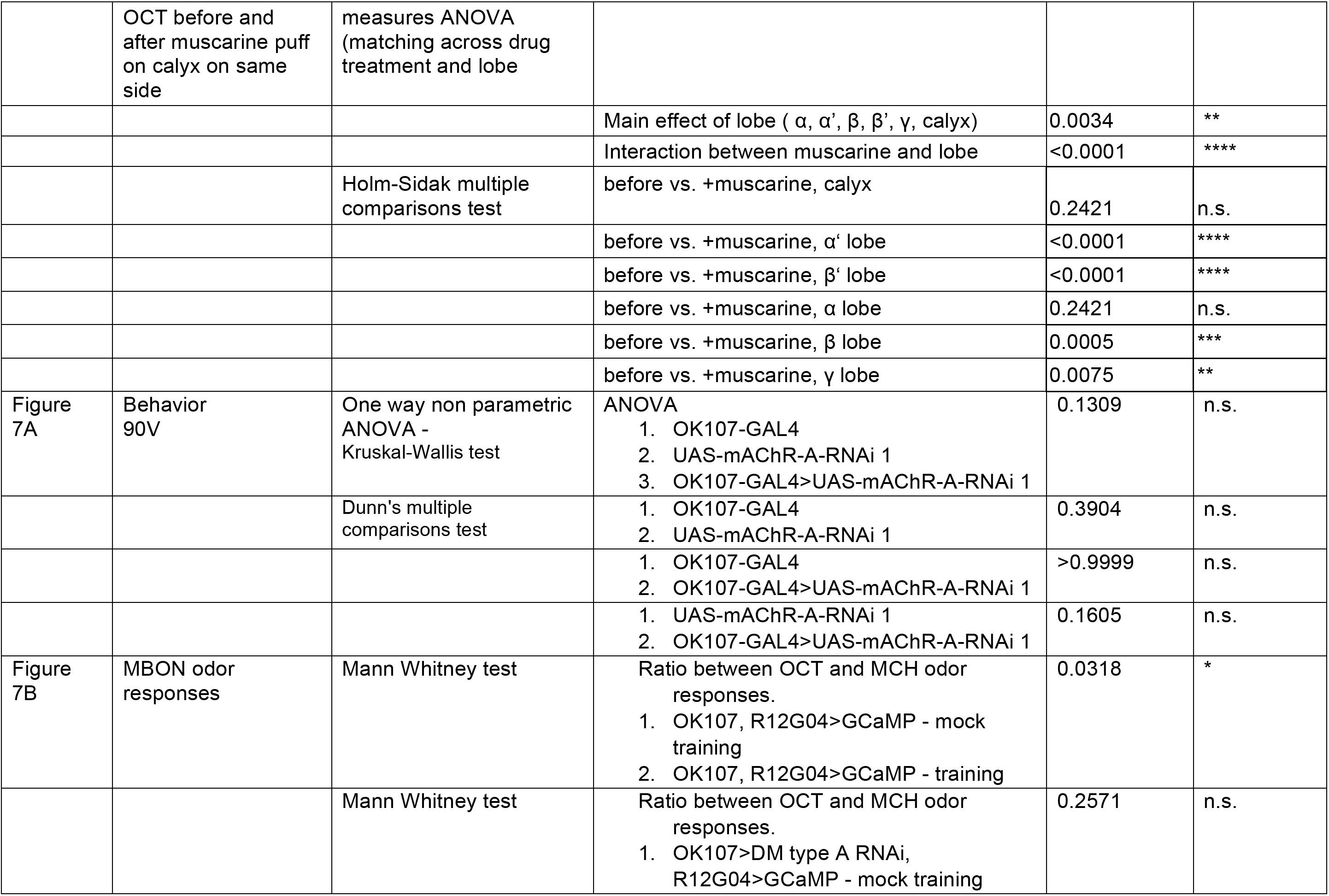
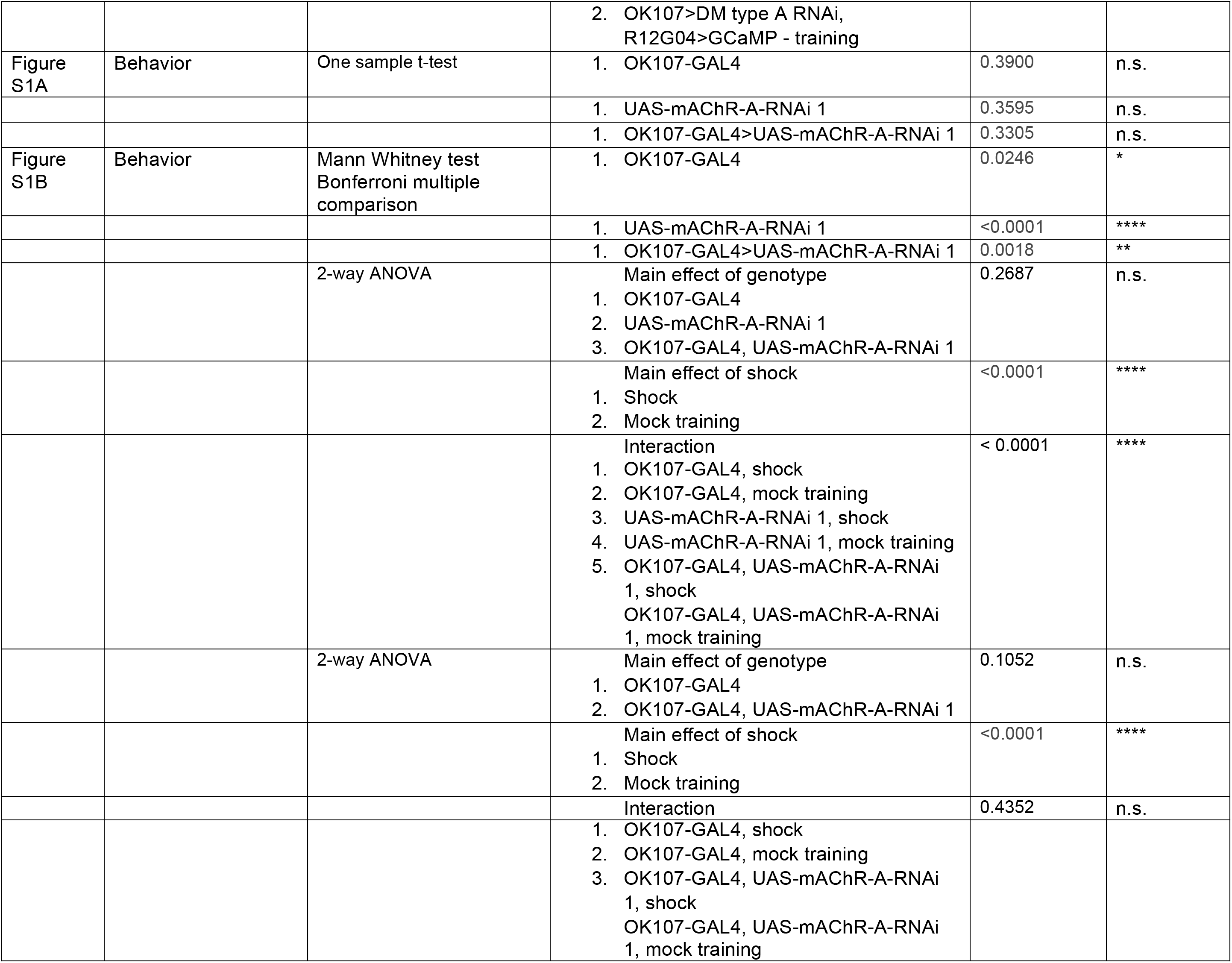
Details of statistical analysis.

**Table S2.** Detailed genotypes used in this study.

| Figure | Shorthand | Full genotype |
| --- | --- | --- |
| 1A, S1 | N/A | *elav-GAL4/+* or *Y*<br>*UAS-mAChR-A-RNAi 1/+*<br>*elav-GAL4/+* or *Y*; +; *UAS-mAChR-A-RNAi 1/+*<br>*UAS-dcr2/+*; *UAS-mAChR-A-RNAi 2/+*<br>*elav-GAL4/+* or *Y*; *UAS-dcr2/+*; *UAS-mAChR-A-RNAi 2/+* |
| 1B,C | N/A | *OK107-GAL4/+*<br>*UAS-mAChR-A-RNAi 1/+*<br>*UAS-mAChR-A-RNAi 1/+*; +; *OK107-GAL4/+*<br>*UAS-dcr2/+*; *UAS-mAChR-A-RNAi 2/+*<br>*UAS-dcr2/+*; *UAS-mAChR-A-RNAi 2/+*; *OK107-GAL4/+* |
| 1E | N/A | *tub-GAL80<sup>ts</sup>/+*; +; *OK107-GAL4/+*<br>*UAS-mAChR-A-RNAi 1/+*<br>*tub-GAL80<sup>ts</sup>/+*; *UAS-mAChR-A-RNAi 1/+*; *OK107-GAL4/+* |
| 2A | N/A | *MiMIC-mAChR-A-GAL4*; *20xUAS-6xGFP* |
| 2B | N/A | *UAS-mAChR-A-RNAi 1/+*<br>*mb247-GAL4/+*<br>*mb247-GAL4/UAS-mAChR-A-RNAi 1*<br>*c305a-GAL4/+*<br>*c305a-GAL4/+*; *UAS-mAChR-A-RNAi 1/+*<br>{*lexAop-GAL80*}*su(Hw)attP5/R45H04-LexA*; *mb247-GAL4/+*<br>{*lexAop-GAL80*}*su(Hw)attP5/R45H04-LexA*; *mb247-GAL4/UAS-mAChR-A-RNAi 1*<br>{*lexAop-GAL80*}*su(Hw)attP5/R44E04-LexA*; *mb247-GAL4/+*<br>{*lexAop-GAL80*}*su(Hw)attP5/R44E04-LexA*; *mb247-GAL4/UAS-mAChR-A-RNAi 1* |
| 3,4 | *OK107-GAL4>GCaMP6f, dcr2* | {*UAS-GCaMP6f*}*attP40/UAS-dcr2*; +; *OK107-GAL4/+* |
| 3,4 | *OK107-GAL4>GCaMP6f, dcr2, mAChR-A-RNAi 2* | {*UAS-GCaMP6f*}*attP40/UAS-dcr2*; *UAS-mAChR-A-RNAi 2/+*; *OK107-GAL4/+* |
| 4 | *mb247-GAL4>GCaMP6f, R44E04-LexA>Gal80* | {*lexAop-GAL80*}*su(Hw)attP5*, {*UAS-GCaMP6f*}*attP40/R44E04-LexA*; *mb247-GAL4/+* |
| 4 | *mb247-GAL4>GCaMP6f, mAChR-A-RNAi 1, R44E04-LexA>Gal80* | {*lexAop-GAL80*}*su(Hw)attP5*, {*UAS-GCaMP6f*}*attP40/R44E04-LexA*; *mb247-GAL4/UAS-mAChR-A-RNAi 1* |
| 5A,B, 6, S4 | OK107-GAL4>GCaMP6f | {UAS-GCaMP6f}attP40/+; +; OK107-GAL4/+ |
| 5C | GH146-GAL4>GCaMP6f | GH146-GAL4/+; UAS-GCaMP6f/+ |
| 5D, S4 | APL unlabeled, APL>TNT | NP2631-GAL4, GH146-FLP/tub-FRT-GAL80-FRT, UAS-mCherry, UAS-TNT; mb247-LexA, lexAop-GCaMP6f/+ |
| 7 | OK107-GAL4, R12G04-LexA>GCaMP6f, mb247-dsRed | {R12G04-LexA}attP40/{lexAop-GCaMP6f}su(Hw)attP5, mb247-dsRed; +; OK107-GAL4/+ |
| 7 | OK107-GAL4>mAChR-A-RNAi 1, R12G04-LexA>GCaMP6f, mb247-dsRed | {R12G04-LexA}attP40/{lexAop-GCaMP6f}su(Hw)attP5, mb247-dsRed; UAS-mAChR-A-RNAi 1/+; OK107-GAL4/+ |
| S3 | OK107-GAL4>UAS-GFP | +; UAS-mCD8-GFP/+;OK107-GAL4/+ |
| S3 | MB247-GAL4>UAS-GFP | +; UAS-mCD8-GFP/mb247-GAL4; + |
| S3 | c305a-GAL4>UAS-GFP | c305a-GAL4, UAS-mCD8-GFP/CyO; Sb/TM3,Ser |
| S3 | R44E04-LexA>LexAop-GFP | {R44E04-LexA}attP40/{LexAop2-mCD8-GFP}attP40/CyO; +; + |
| S3 | R45H04-LexA>LexAop-GFP | {R45H04-LexA}attP40/{LexAop2-mCD8-GFP}attP40/CyO; +; + |
| S3 | MB247-GAL4>UAS-GFP<br>R45H04-LexA>LexAop-GAL80 | {R45H04-LexA}attP40/UAS-mCD8-GFP; mb247-GAL4/LexAop-GAL80; + |
| S3 | MB247-GAL4> UAS-GCaMP6f<br>R44E04-LexA>LexAop-GAL80 | {R44E04-LexA}attP40/LexAop-GAL80, UAS-GCaMP6f; mb247-GAL4/+ |
| S3 | R12G04-LexA>LexAop-GFP | {R12G04-LexA}attP40/LexAop2-mCD8-GFP; +; + |

## References

Allen, T.G., and Brown, D.A. (1993). M2 muscarinic receptor-mediated inhibition of the Ca2+ current in rat magnocellular cholinergic basal forebrain neurones. J. Physiol. (Lond.) 466, 173–189.

Anagnostaras, S.G., Murphy, G.G., Hamilton, S.E., Mitchell, S.L., Rahnama, N.P., Nathanson, N.M., and Silva, A.J. (2003). Selective cognitive dysfunction in acetylcholine M1 muscarinic receptor mutant mice. Nat Neurosci 6, 51–58.

Aso, Y., and Rubin, G.M. (2016). Dopaminergic neurons write and update memories with cell-type-specific rules. Elife 5, e16135.

Aso, Y., Hattori, D., Yu, Y., Johnston, R.M., Iyer, N.A., Ngo, T.-T.B., Dionne, H., Abbott, L.F., Axel, R., Tanimoto, H., et al. (2014a). The neuronal architecture of the mushroom body provides a logic for associative learning. Elife 3, e04577.

Aso, Y., Sitaraman, D., Ichinose, T., Kaun, K.R., Vogt, K., Belliart-Guérin, G., Plaçais, P.-Y., Robie, A.A., Yamagata, N., Schnaitmann, C., et al. (2014b). Mushroom body output neurons encode valence and guide memory-based action selection in Drosophila. Elife 3, e04580.

Barnstedt, O., Owald, D., Felsenberg, J., Brain, R., Moszynski, J.-P., Talbot, C.B., Perrat, P.N., and Waddell, S. (2016). Memory-Relevant Mushroom Body Output Synapses Are Cholinergic. Neuron 89, 1237–1247.

Becnel, J., Johnson, O., Majeed, Z.R., Tran, V., Yu, B., Roth, B.L., Cooper, R.L., Kerut, E.K., and Nichols, C.D. (2013). DREADDs in Drosophila: a pharmacogenetic approach for controlling behavior, neuronal signaling, and physiology in the fly. Cell Rep 4, 1049–1059.

Bellingham, M.C., and Berger, A.J. (1996). Presynaptic depression of excitatory synaptic inputs to rat hypoglossal motoneurons by muscarinic M2 receptors. J. Neurophysiol. 76, 3758–3770.

Berry, J.A., Cervantes-Sandoval, I., Nicholas, E.P., and Davis, R.L. (2012). Dopamine Is Required for Learning and Forgetting in Drosophila. Neuron 74, 530–542.

Blake, A.D., Anthony, N.M., Chen, H.H., Harrison, J.B., Nathanson, N.M., and Sattelle, D.B. (1993). Drosophila nervous system muscarinic acetylcholine receptor: transient functional expression and localization by immunocytochemistry. Mol. Pharmacol. 44, 716–724.

Bräcker, L.B., Siju, K.P., Varela, N., Aso, Y., Zhang, M., Hein, I., Vasconcelos, M.L., and Kadow, I.C.G. (2013). Essential Role of the Mushroom Body in Context-Dependent CO2 Avoidance in Drosophila. Curr. Biol. 1–7.

Busto, G.U., Cervantes-Sandoval, I., and Davis, R.L. (2010). Olfactory learning in Drosophila. Physiology 25, 338–346.

Campbell, R.A.A., Honegger, K.S., Qin, H., Li, W., Demir, E., and Turner, G.C. (2013). Imaging a population code for odor identity in the Drosophila mushroom body. J. Neurosci. 33, 10568–10581.

Cantrell, A.R., Ma, J.Y., Scheuer, T., and Catterall, W.A. (1996). Muscarinic modulation of sodium current by activation of protein kinase C in rat hippocampal neurons. Neuron 16, 1019–1026.

Caulfield, M.P., and Birdsall, N.J.M. (1998). International Union of Pharmacology. XVII. Classification of Muscarinic Acetylcholine Receptors. Pharmacol. Rev. 50, 279–290.

Cervantes-Sandoval, I., Phan, A., Chakraborty, M., and Davis, R.L. (2017). Reciprocal synapses between mushroom body and dopamine neurons form a positive feedback loop required for learning. Elife 6.

Chen, T.-W., Wardill, T.J., Sun, Y., Pulver, S.R., Renninger, S.L., Baohan, A., Schreiter, E.R., Kerr, R.A., Orger, M.B., Jayaraman, V., et al. (2013). Ultrasensitive fluorescent proteins for imaging neuronal activity. Nature 499, 295–300.

Christiansen, F., Zube, C., Andlauer, T.F.M., Wichmann, C., Fouquet, W., Owald, D., Mertel, S., Leiss, F., Tavosanis, G., Farca Luna, A.J., et al. (2011). Presynapses in Kenyon Cell Dendrites in the Mushroom Body Calyx of Drosophila. Journal of Neuroscience 31, 9696–9707.

Claridge-Chang, A., Roorda, R.D., Vrontou, E., Sjulson, L., Li, H., Hirsh, J., and Miesenböck, G. (2009). Writing memories with light-addressable reinforcement circuitry. Cell 139, 405–415.

Cognigni, P., Felsenberg, J., and Waddell, S. (2017). Do the right thing: neural network mechanisms of memory formation, expression and update in Drosophila. Curr Opin Neurobiol 49, 51–58.

Cohn, R., Morantte, I., and Ruta, V. (2015). Coordinated and Compartmentalized Neuromodulation Shapes Sensory Processing in Drosophila. Cell 163, 1742–1755.

Collin, C., Hauser, F., Gonzalez de Valdivia, E., de Valdivia, E.G., Li, S., Reisenberger, J., Carlsen, E.M.M., Khan, Z., Hansen, N.O., Puhm, F., et al. (2013). Two types of muscarinic acetylcholine receptors in Drosophila and other arthropods. Cell. Mol. Life Sci. 70, 3231–3242.

Connolly, J.B., Roberts, I.J., Armstrong, J.D., Kaiser, K., Forte, M., Tully, T., and O’Kane, C.J. (1996). Associative learning disrupted by impaired Gs signaling in Drosophila mushroom bodies. Science 274, 2104–2107.

Croset, V., Treiber, C.D., and Waddell, S. (2018). Cellular diversity in the Drosophila midbrain revealed by single-cell transcriptomics. Elife 7, e34550.

Davie, K., Janssens, J., Koldere, D., De Waegeneer, M., Pech, U., Kreft, Ł., Aibar, S., Makhzami, S., Christiaens, V., Bravo González-Blas, C., et al. (2018). A Single-Cell Transcriptome Atlas of the Aging Drosophila Brain. Cell.

de Vin, F., Choi, S.M., Bolognesi, M.L., and Lefebvre, R.A. (2015). Presynaptic M 3 muscarinic cholinoceptors mediate inhibition of excitatory synaptic transmission in area CA1 of rat hippocampus. Brain Research.

Dubbs, A., Guevara, J., and Yuste, R. (2016). moco: Fast Motion Correction for Calcium Imaging. Front Neuroinform 10, 6.

Eichler, K., Li, F., Litwin-Kumar, A., Park, Y., Andrade, I., Schneider-Mizell, C.M., Saumweber, T., Huser, A., Eschbach, C., Gerber, B., et al. (2017). The complete connectome of a learning and memory centre in an insect brain. Nature 548, 175–182.

Farris, S.M. (2011). Are mushroom bodies cerebellum-like structures? Arthropod Structure and Development 40, 368–379.

Gamper, N., Reznikov, V., Yamada, Y., Yang, J., and Shapiro, M.S. (2004). Phosphatidylinositol [correction] 4,5-bisphosphate signals underlie receptor-specific Gq/11-mediated modulation of N-type Ca2+ channels. J. Neurosci. 24, 10980–10992.

Groschner, L.N., Chan Wah Hak, L., Bogacz, R., DasGupta, S., and Miesenböck, G. (2018). Dendritic Integration of Sensory Evidence in Perceptual Decision-Making. Cell 173, 894–905.e13.

Guven-Ozkan, T., and Davis, R.L. (2014). Functional neuroanatomy of Drosophila olfactory memory formation. Learning & Memory 21, 519–526.

Hannan, F., and Hall, L.M. (1996). Temporal and spatial expression patterns of two G-protein coupled receptors inDrosophila melanogaster. Invertebrate Neuroscience.

Haynes, P.R., Christmann, B.L., and Griffith, L.C. (2015). A single pair of neurons links sleep to memory consolidation in Drosophila melanogaster. Elife 4.

Hige, T., Aso, Y., Rubin, G.M., and Turner, G.C. (2015a). Plasticity-driven individualization of olfactory coding in mushroom body output neurons. Nature.

Hige, T. (2017). What can tiny mushrooms in fruit flies tell us about learning and memory? Neurosci. Res.

Hige, T., Aso, Y., Modi, M.N., Rubin, G.M., and Turner, G.C. (2015b). Heterosynaptic Plasticity Underlies Aversive Olfactory Learning in Drosophila. Neuron 88, 985–998.

Himmelreich, S., Masuho, I., Berry, J.A., MacMullen, C., Skamangas, N.K., Martemyanov, K.A., and Davis, R.L. (2017). Dopamine Receptor DAMB Signals via Gq to Mediate Forgetting in Drosophila. Cell Rep 21, 2074–2081.

Jenett, A., Rubin, G.M., Ngo, T.-T.B., Shepherd, D., Murphy, C., Dionne, H., Pfeiffer, B.D., Cavallaro, A., Hall, D., Jeter, J., et al. (2012). A GAL4-driver line resource for Drosophila neurobiology. Cell Rep 2, 991–1001.

Jiang, L., Kosenko, A., Yu, C., Huang, L., Li, X., and Hoshi, N. (2015). Activation of m1 muscarinic acetylcholine receptor induces surface transport of KCNQ channels through a CRMP-2-mediated pathway. J. Cell. Sci. 128, 4235–4245.

Jo, J., Son, G.H., Winters, B.L., Kim, M.J., Whitcomb, D.J., Dickinson, B.A., Lee, Y.-B., Futai, K., Amici, M., Sheng, M., et al. (2010). Muscarinic receptors induce LTD of NMDAR EPSCs via a mechanism involving hippocalcin, AP2 and PSD-95. Nat Neurosci 13, 1216–1224.

Jörntell, H., and Hansel, C. (2006). Synaptic memories upside down: bidirectional plasticity at cerebellar parallel fiber-Purkinje cell synapses. Neuron 52, 227–238.

Kammermeier, P.J., Ruiz-Velasco, V., and Ikeda, S.R. (2000). A voltage-independent calcium current inhibitory pathway activated by muscarinic agonists in rat sympathetic neurons requires both Galpha q/11 and Gbeta gamma. Journal of Neuroscience 20, 5623–5629.

Kamsler, A., McHugh, T.J., Gerber, D., Huang, S.Y., and Tonegawa, S. (2010). Presynaptic m1 muscarinic receptors are necessary for mGluR long-term depression in the hippocampus. Proc. Natl. Acad. Sci. USA 107, 1618–1623.

Keum, D., Baek, C., Kim, D.-I., Kweon, H.-J., and Suh, B.-C. (2014). Voltage-dependent regulation of CaV2.2 channels by Gq-coupled receptor is facilitated by membrane-localized β subunit. J. Gen. Physiol. 144, 297–309.

Krashes, M.J., Keene, A.C., Leung, B., Armstrong, J.D., and Waddell, S. (2007). Sequential Use of Mushroom Body Neuron Subsets during Drosophila Odor Memory Processing. Neuron 53, 103–115.

Lei, Z., Chen, K., Li, H., Liu, H., and Guo, A. (2013). The GABA system regulates the sparse coding of odors in the mushroom bodies of Drosophila. Biochem. Biophys. Res. Commun. 436, 35–40.

Leitch, B., and Laurent, G. (1996). GABAergic synapses in the antennal lobe and mushroom body of the locust olfactory system. J. Comp. Neurol.

Lin, A.C., Bygrave, A.M., de Calignon, A., Lee, T., and Miesenböck, G. (2014). Sparse, decorrelated odor coding in the mushroom body enhances learned odor discrimination. Nat Neurosci 17, 559–568.

Lin, D.M., and Goodman, C.S. (1994). Ectopic and increased expression of Fasciclin II alters motoneuron growth cone guidance. Neuron 13, 507–523.

Liu, X., and Davis, R.L. (2008). The GABAergic anterior paired lateral neuron suppresses and is suppressed by olfactory learning. Nat Neurosci 12, 53–59.

Lüscher, C., and Huber, K.M. (2010). Group 1 mGluR-dependent synaptic long-term depression: mechanisms and implications for circuitry and disease. Neuron 65, 445–459.

Masuda-Nakagawa, L.M., Ito, K., Awasaki, T., and O’Kane, C.J. (2014). A single GABAergic neuron mediates feedback of odor-evoked signals in the mushroom body of larval Drosophila. Front. Neural. Circuits 8.

McGuire, S.E., Le, P.T., Osborn, A.J., Matsumoto, K., and Davis, R.L. (2003). Spatiotemporal rescue of memory dysfunction in Drosophila. Science 302, 1765–1768.

Ng, M.M., Roorda, R.D.R., Lima, S.Q., Boris V BV Zemelman, Morcillo, P.P., and Miesenböck, G. (2002). Transmission of olfactory information between three populations of neurons in the antennal lobe of the fly. Neuron 36, 463–474.

Owald, D., Felsenberg, J., Talbot, C.B., Das, G., Perisse, E., Huetteroth, W., and Waddell, S. (2015). Activity of defined mushroom body output neurons underlies learned olfactory behavior in Drosophila. Neuron 86, 417–427.

Papadopoulou, M., Cassenaer, S., Nowotny, T., and Laurent, G. (2011). Normalization for sparse encoding of odors by a wide-field interneuron. Science 332, 721–725.

Parnas, M., Lin, A.C., Huetteroth, W., and Miesenböck, G. (2013). Odor discrimination in Drosophila: from neural population codes to behavior. Neuron 79, 932–944.

Perisse, E., Owald, D., Barnstedt, O., Talbot, C.B., Huetteroth, W., and Waddell, S. (2016). Aversive Learning and Appetitive Motivation Toggle Feed-Forward Inhibition in the Drosophila Mushroom Body. Neuron 90, 1086–1099.

Perisse, E., Yin, Y., Lin, A.C., Lin, S., Huetteroth, W., and Waddell, S. (2013). Different Kenyon cell populations drive learned approach and avoidance in Drosophila. Neuron 79, 945–956.

Pinheiro, P.S., and Mulle, C. (2008). Presynaptic glutamate receptors: physiological functions and mechanisms of action. Nat Rev Neurosci 9, 423–436.

Pitman, J.L., Huetteroth, W., Burke, C.J., Krashes, M.J., Lai, S.-L., Lee, T., and Waddell, S. (2011). A pair of inhibitory neurons are required to sustain labile memory in the Drosophila mushroom body. Curr. Biol. 21, 855–861.

Podgorski, K., Terpetschnig, E., Klochko, O.P., Obukhova, O.M., and Haas, K. (2012). Ultra-bright and -stable red and near-infrared squaraine fluorophores for in vivo two-photon imaging. PLoS ONE 7, e51980.

Qin, H., Cressy, M., Li, W., Coravos, J.S., Izzi, S.A., and Dubnau, J. (2012). Gamma Neurons Mediate Dopaminergic Input during Aversive Olfactory Memory Formation in Drosophila. Curr. Biol. 1–7.

Ren, G.R., Folke, J., Hauser, F., Li, S., and Grimmelikhuijzen, C.J.P. (2015). The A- and B-type muscarinic acetylcholine receptors from Drosophila melanogaster couple to different second messenger pathways. Biochem. Biophys. Res. Commun. 462, 358–364.

Riemensperger, T., Völler, T., Stock, P., Buchner, E., and Fiala, A. (2005). Punishment prediction by dopaminergic neurons in Drosophila. Curr. Biol. 15, 1953–1960.

Scanziani, M., Salin, P.A., Vogt, K.E., Malenka, R.C., and Nicoll, R.A. (1997). Use-dependent increases in glutamate concentration activate presynaptic metabotropic glutamate receptors. Nature 385, 630–634.

Schonewille, M., Gao, Z., Boele, H.-J., Veloz, M.F.V., Amerika, W.E., Simek, A.A.M., De Jeu, M.T., Steinberg, J.P., Takamiya, K., Hoebeek, F.E., et al. (2011). Reevaluating the role of LTD in cerebellar motor learning. Neuron 70, 43–50.

Schürmann, F.-W. (2016). Fine structure of synaptic sites and circuits in mushroom bodies of insect brains. Arthropod Structure and Development 45, 399–421.

Selcho, M., Pauls, D., Han, K.-A., Stocker, R.F., and Thum, A.S. (2009). The role of dopamine in Drosophila larval classical olfactory conditioning. PLoS ONE 4, e5897.

Séjourné, J., Plaçais, P.-Y., Aso, Y., Siwanowicz, I., Trannoy, S., Thoma, V., Tedjakumala, S.R., Rubin, G.M., Tchénio, P., Ito, K., et al. (2011). Mushroom body efferent neurons responsible for aversive olfactory memory retrieval in Drosophila. Nat Neurosci 14, 903–910.

Shearin, H.K., Macdonald, I.S., Spector, L.P., and Stowers, R.S. (2014). Hexameric GFP and mCherry reporters for the Drosophila GAL4, Q, and LexA transcription systems. Genetics 196, 951–960.

Sheridan, R.D., and Sutor, B. (1990). Presynaptic M 1 muscarinic cholinoceptors mediate inhibition of excitatory synaptic transmission in the hippocampus in vitro. Neurosci. Lett.

Silva, B., Molina-Fernández, C., Ugalde, M.B., Tognarelli, E.I., Angel, C., and Campusano, J.M. (2015). Muscarinic ACh Receptors Contribute to Aversive Olfactory Learning in Drosophila. Neural Plast. 2015, 1–10.

Slutsky, I., Parnas, H., and Parnas, I. (1999). Presynaptic effects of muscarine on ACh release at the frog neuromuscular junction. J. Physiol. (Lond.) 514 (Pt 3), 769–782.

Stocker, R.F., Heimbeck, G., Gendre, N., and de Belle, J.S. (1997). Neuroblast ablation in Drosophila P[GAL4] lines reveals origins of olfactory interneurons. J. Neurobiol. 32, 443–456.

Strausfeld, N.J., and Li, Y. (1999). Representation of the calyces in the medial and vertical lobes of cockroach mushroom bodies. J. Comp. Neurol. 409, 626–646.

Su, H., and O’Dowd, D.K. (2003). Fast synaptic currents in Drosophila mushroom body Kenyon cells are mediated by alpha-bungarotoxin-sensitive nicotinic acetylcholine receptors and picrotoxin-sensitive GABA receptors. Journal of Neuroscience 23, 9246–9253.

Suh, B.-C., Leal, K., and Hille, B. (2010). Modulation of high-voltage activated Ca(2+) channels by membrane phosphatidylinositol 4,5-bisphosphate. Neuron 67, 224–238.

Takemura, S.-Y., Aso, Y., Hige, T., Wong, A., Lu, Z., Xu, C.S., Rivlin, P.K., Hess, H., Zhao, T., Parag, T., et al. (2017). A connectome of a learning and memory center in the adult Drosophila brain. Elife 6.

Tully, T., and Quinn, W.G. (1985). Classical conditioning and retention in normal and mutant Drosophila melanogaster. J. Comp. Physiol. (a) 157, 263–277.

Turner, G.C., Bazhenov, M., and Laurent, G. (2008). Olfactory representations by Drosophila mushroom body neurons. J. Neurophysiol. 99, 734–746.

Venken, K.J.T., Schulze, K.L., Haelterman, N.A., Pan, H., He, Y., Evans-Holm, M., Carlson, J.W., Levis, R.W., Spradling, A.C., Hoskins, R.A., et al. (2011). MiMIC: a highly versatile transposon insertion resource for engineering Drosophila melanogaster genes. Nat Methods 8, 737–743.

Volk, L.J., Pfeiffer, B.E., Gibson, J.R., and Huber, K.M. (2007). Multiple Gq-coupled receptors converge on a common protein synthesis-dependent long-term depression that is affected in fragile X syndrome mental retardation. J. Neurosci. 27, 11624–11634.

Wang, J.W., Wong, A.M., Flores, J., Vosshall, L.B., and Axel, R. (2003). Two-photon calcium imaging reveals an odor-evoked map of activity in the fly brain. Cell 112, 271–282.

Wu, C.-L., Shih, M.-F.M., Lee, P.-T., and Chiang, A.-S. (2013). An octopamine-mushroom body circuit modulates the formation of anesthesia-resistant memory in Drosophila. Curr. Biol. 23, 2346–2354.

Wu, J.S., and Luo, L. (2006). A protocol for dissecting Drosophila melanogaster brains for live imaging or immunostaining. Nat Protoc 1, 2110–2115.

Yamaguchi, K., Itohara, S., and Ito, M. (2016). Reassessment of long-term depression in cerebellar Purkinje cells in mice carrying mutated GluA2 C terminus. Proc. Natl. Acad. Sci. USA 113, 10192–10197.

Yasuyama, K., and Salvaterra, P.M. (1999). Localization of choline acetyltransferase-expressing neurons in Drosophila nervous system. Microsc. Res. Tech. 45, 65–79.

Zars, T. (2000). Localization of a Short-Term Memory in Drosophila. Science 288, 672–675.

